# How does plant chemodiversity evolve? Testing five hypotheses in one population genetic model

**DOI:** 10.1101/2024.04.12.589236

**Authors:** Meike J. Wittmann, Andrea Bräutigam

## Abstract

- Plant chemodiversity, the diversity of plant specialized metabolites, is an important dimension of biodiversity. However, there are so far few quantitative models to test verbal hypotheses on how chemodiversity evolved. Here we develop such a model to test predictions of five hypotheses: the “fluctuating selection hypothesis”, the “dominance reversal hypothesis”, the interaction diversity hypothesis, the synergy hypothesis, and the screening hypothesis.
- We build a population genetic model of a plant population attacked by herbivore species whose occurrence fluctuates over time. We study the model using mathematical analysis and individual-based simulations.
- As predicted by the “dominance reversal hypothesis”, chemodiversity can be maintained if alleles conferring a defense metabolite are dominant with respect to the benefits, but recessive with respect to costs. However, even smaller changes in dominance can maintain polymorphism. Moreover, our results underpin and elaborate predictions of the synergy and interaction diversity hypotheses, and, to the extent that our model can address it, the screening hypotheses. By contrast, we found only partial support for the “fluctuating selection hypothesis”.
- In summary, we have developed a flexible model and tested various verbal models for the evolution of chemodiversity. Next, more mechanistic models are needed that explicitly consider the organization of metabolic pathways.

## Introduction

Plants harbor an amazing diversity of so-called specialised (also known as secondary) metabolites. These are metabolites that are not involved in basic functions like photosynthesis that are shared by (almost) all plants. Specialised metabolites are usually only found in some plant lineages and are often involved in defense against herbivores or attraction of pollinators. For many metabolites, their function is still unknown. The diversity of specialised metabolites, also known as chemodiversity, is found at many levels of organisations: within and between individuals, populations, communities, species, and larger taxonomic groups (Wetzel and Whitehead 2020). Chemodiversity raises several evolutionary questions: Why are there so many metabolites? And why do individuals differ qualitatively and quantitatively in the metabolites that they produce? That is, what maintains polymorphism in populations with respect to metabolites?

Potential answers to such questions are offered by a number of hypotheses and verbal models (for reviews, see Moore *et al*. 2014; Thon *et al*. 2024; Wetzel and Whitehead 2020). To test whether these verbal models “work”, i.e., whether the claimed predictions actually follow from the assumptions, and to find out what exact patterns of chemodiversity they predict, quantitative models such as simple mathematical models and detailed computer simulations are needed (Servedio *et al*. 2014). Such models exist for genetic variation and for species diversity, but so far there is a relatively small number of quantitative models for chemodiversity (reviewed in Thon *et al*. 2024). The existing quantitative models include optimality models that predict investment in defenses based on resource constraints (e.g., Coley *et al*. 1985; Orrock *et al*. 2015; Yamamura and Tsuji 1995), evolutionary game theory and frequency-dependent selection models that explain coexistence between defended and undefended plants based on direct and indirect interactions between plants in neighborhoods (e.g., Augner *et al*. 1991; Lankau 2009; Sato *et al*. 2017), as well as simulation models for plant-herbivore coevolution (e.g., Speed *et al*. 2015). While some of these models specifically address variation in chemical traits, others more generally address plant defense traits, but could be also be applied to chemical traits.

However, several important verbal chemodiversity hypotheses that are often used to generate hypotheses for empirical work so far lack underpinning by quantitative models. First, the interaction diversity hypothesis (Iason *et al*. 2011; Whitehead *et al*. 2021) posits that plants produce many different metabolites because they are engaged in many different ecological interactions that each are mediated by different metabolites. Second, the synergy hypothesis suggests that if there are synergistic benefits of defense metabolites, where the anti-herbivore effects of mixtures are larger than expected based on adding the effects of individual metabolites, we expect plants to produce a larger number of such metabolites (Richards *et al*. 2016). Third, the screening hypothesis (Firn and Jones 2003; Jones and Firn 1991) suggests that each new metabolite has only a small probability of having biological activity in a relevant ecological interaction, e.g. against a herbivore and thus predicts that plants need to produce a large number of metabolites to “find” the few useful ones. Since the machinery for producing metabolites is costly, the screening hypothesis further predicts that plants should evolve grid-like pathways with promiscuous enzymes that can produce a large diversity of metabolites very efficiently.

Furthermore, although much of chemodiversity has a strong genetic basis and will be strongly affected also by processes like mutation and genetic drift, there is so far a surprising lack of population genetic models for chemodiversity. Based on models for other types of genetic variation (see e.g. Haldane and Jayakar 1963; Hedrick 1976; Johnson *et al*. 2023; Wittmann *et al*. 2017), temporally fluctuating selection can be a powerful mechanism for the maintenance of genetic polymorphism, and thus potentially for differences between individuals in the metabolites that they produce. Since many insect herbivores fluctuate in abundance over time (De-la-Cruz and Núñez-Farfán 2023; Root 1996; Stange *et al*. 2011), e.g. due to temperature variation between seasons or across years, fluctuating selection could be an important mechanism for the maintenance of chemodiversity between individuals in a population. For example, if there are trade-offs where metabolites repel generalist herbivores, but may also attract specialist herbivores, genetic polymorphism underlying chemodiversity could potentially be maintained by fluctuations in the presence of specialist and generalist herbivores (as suggested for the glucosinolate sinigrin by Lankau 2007). However, this “fluctuating-selection hypothesis” has not been explored in a quantitative model.

Finally, we know from population genetic theory that whether or not polymorphism is maintained can strongly depend on patterns of genetic dominance. Dominance quantifies the phenotype or fitness of heterozygotes relative to homozygotes. Consider, for example, a locus with two alleles, 1 (presence) and 0 (absence), where “11” individuals produce a metabolite, but “00” individuals do not. Then the 1 allele would be called (partially) dominant if “10” heterozygotes produce more than half of what 11 homozygotes produce and recessive if heterozygotes produce less than half. In particular, reversals of dominance across contexts can be a powerful mechanism for the maintenance of genetic variation (Connallon and Chenoweth 2019; Curtsinger *et al*. 1994; Grieshop *et al*. 2024; Hoekstra *et al*. 1985; Rose 1982; Wittmann *et al*. 2017). For example, polymorphism could be maintained if the 1 allele promotes viability and is also dominant for viability, but reduces fecundity and is recessive for fecundity. So far there is no modeling work yet exploring this “dominance-reversal hypothesis” for chemodiversity. From crossing experiments we know that spe-cialised metabolites and plant defense traits often exhibit dominance (Orians 2000; van Dam and Baldwin 2003), although to our knowledge nothing is known on dominance reversals.

To help fill these gaps in chemodiversity modeling, in this paper we therefore develop a population genetic modeling approach focused on the evolution of chemodiversity via presence-absence polymorphisms, i.e., polymorphisms where some individuals in the population produce a metabolite while others do not. We use this model to explore how patterns of chemodiversity are affected by temporal fluctuations in herbivore occurrence, dominance patterns, the number of herbivores, and synergistic effects of metabolites. For each scenario, we will analytically (if possible) determine the potential for genetic polymorphism and use individual-based simulations to quantify the expected total number of metabolites produced in the population (*γ* diversity), the average number of metabolites produced per individual (*α* diversity), and the average number of different metabolites between two individuals (*β* diversity). Thereby we are able to perform proof-of-concept tests for the dominance reversal hypothesis, the fluctuating selection hypothesis, the interaction diversity hypothesis and the synergy hypothesis. Additionally, we test whether plant populations with higher chemodiversity are better able to adapt to the invasion of a new herbivore, a conjecture that can be derived from the screening hypothesis.

## Description

We assume a diploid, randomly mating plant population of constant size *N* (see Table 1 for an overview of all parameters and their default values). We focus on the case of annual plants without a seed bank, that is, with discrete non-overlapping generations, but also consider the case with overlapping generations with an adult death probability *θ* per time step (but also without a seed bank). Setting *θ* to 1 corresponds to the case with non-overlapping generations. The plant population can be attacked by *n*_h_ herbivores. Time is divided into phases of *g* generations, where each phase could for example represent a season or the yearly growing season. We assume that herbivore *i* is present and active with probability *p*_*i*_ independently for each phase and independently from other herbivores.

**Table 1.** Parameters and their default values. Unless noted otherwise, the parameters are kept at their default values.

| Parameter | Explanation | Default value |
| --- | --- | --- |
| $N$ | Population size | 500 |
| $\theta$ | Adult death probability per time step | 1 |
| $n_h$ | Number of herbivores | 1 |
| $g$ | Number of generations per phase | 5 |
| $p_i$ | Probability that herbivore $i$ is present and active during a particular phase | 0.2 |
| $L$ | Number of loci responsible for metabolite production = maximum possible number of different metabolites | 10 |
| $m_{li}$ | Protective effect of metabolite $l$ against herbivore $i$ | 1 |
| $d_a$ | Dominance for effect on herbivores | always varied |
| $d_c$ | Dominance for costs | always varied |
| $b_0$ | Baseline probability to escape herbivore | 0 |
| $a_{\text{half}}$ | Half-saturation parameter of the benefit function | 0 |
| $s$ | Sensitivity parameter of the benefit function | 1 |
| $c$ | Cost parameter | 0.02 |
| $u$ | Mutation probability per allele copy per generation | 0.0001 |

The plant genome has *L* loci responsible for the production of metabolites that affect herbivores. They could be either the genes coding for the biosynthetic enzymes or regulatory loci affecting such genes. The *L × n*_h_ matrix **M** = (*m*_*li*_) specifies anti-herbivore activity of a certain locus *l* against a certain herbivore *i*. If *m*_*li*_ *>* 0, the metabolite acts as a repellent for herbivore *i*, if *m*_*li*_ = 0 it has no effect, and if *m*_*li*_ *<* 0 the metabolite attracts the herbivore. At each locus, there can be two alleles: 0 (absence, i.e. a deletion or a non-functional gene-variant) and 1 (presence). That is, individuals can have three possible genotypes: 00 or 11 homozygote or 01 heterozygote/hemizygote.

For each individual and for each herbivore *i*, we first determine what anti-herbivore activity *a*_*i*_ the individual has against herbivore *i*. We here assume that activities are additive across loci but that there can be dominance effects:

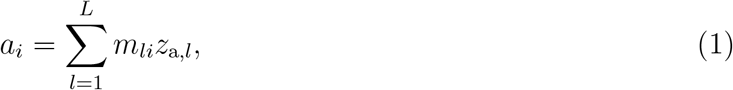

where

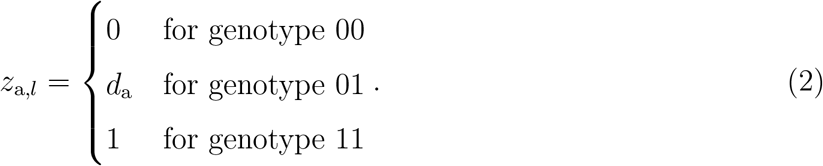

We assume that the “activity dominance” *d*_a_ is between 0 and 1 and for simplicity is constant across loci. Activity dominance 0 means that heterozygotes do not benefit at all from the single copy of the presence allele, whereas an activity dominance of 1 means that they enjoy the same protective effects as an individual with two copies of the presence allele.

The activity *a* against a certain herbivore then affects fitness at times when the herbivore is present. We capture this in a benefit function *b*(*a*). Different shapes of this function are possible.

In the following, we use a classical logistic benefit function

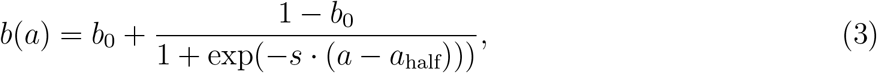

where *b*_0_ is the baseline probability to escape the herbivore, in the worst case that an individual is strongly attracting herbivores (large negative *a*). The parameter *b*_0_ can be used to model different levels of herbivore pressure. *a*_half_ is a half-saturation parameter specifying at which activity level half of the possible additional protection is achieved, and *s* is the sensitivity of the protection to differences in the anti-herbivore activity (Fig. 1 a). The output *b*(*a*) can be understood as the proportion of plant material not consumed by the herbivore or more abstractly as the proportion of fitness remaining after a generation where the respective herbivore is present. We chose this function because the protection from herbivores approaches 1 for very high activity levels and with different parameter choices, in particular *a*_half_, the benefit function can be saturating, at least over most of the positive range of activity levels or exhibit synergy (Fig. 1 b). This enables us to test predictions of the synergy hypothesis.

**Figure 1.**
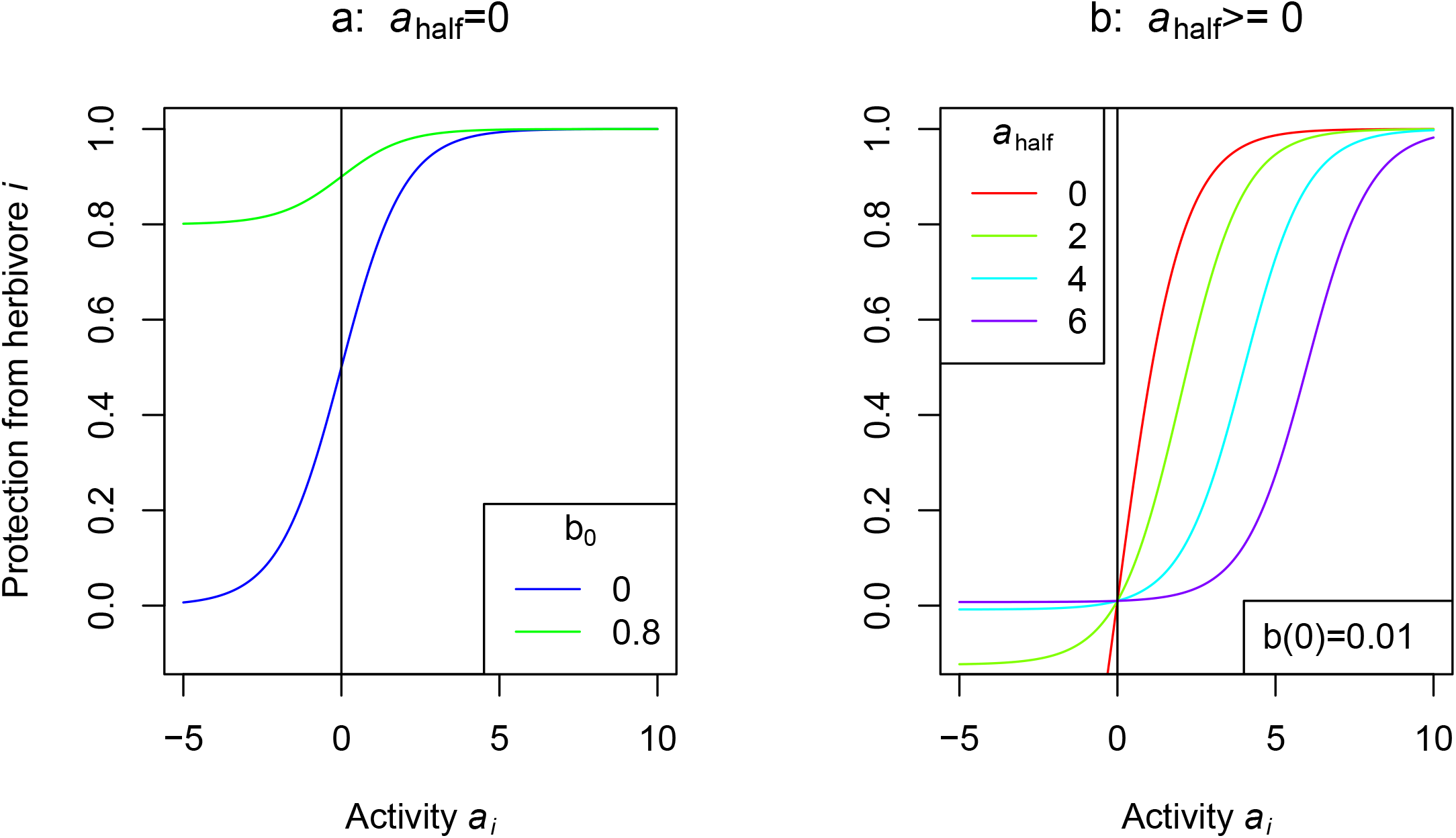
Benefit functions *b*(*a*_*i*_) used in this study. In (a), the green curve represents a low-herbivory scenario, whereas the blue curve represents a high-herbivory scenario. In (b), benefit functions with increasing *a*_half_ exhibit increasing synergy. The baseline probability to escape the herbivore *b*_0_ for each scenario is chosen such that *b*(0) = 0.01 in all four scenarios.

In addition to their protective benefits, metabolites also have costs. We model these by assuming that fecundity is proportional to the cost function

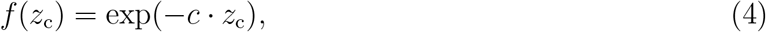

where *c* is a cost parameter and

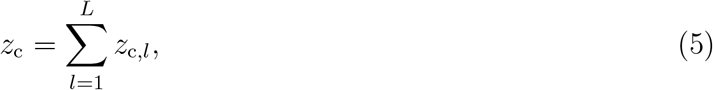

is a “cost trait” with

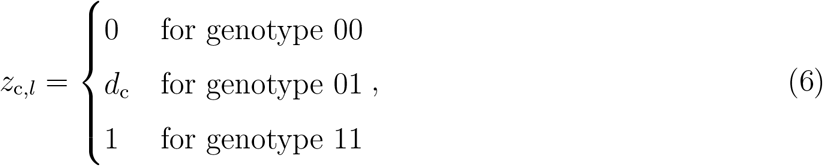

where *d*_c_ is the dominance parameter with respect to costs and is also assumed to be between 0 and 1.

An individual with activities 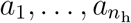 against the various herbivores and “cost trait” *z*_c_ then has the following fitness in generation *t*:

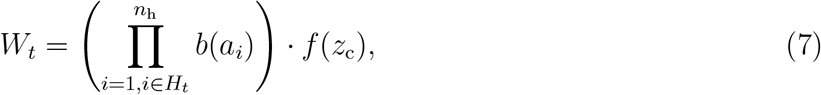

where *H*_*t*_ is the set of herbivores present in generation *t*. That is, we assume that all fitness components act multiplicatively.

Based on the adult death probability *θ*, the number of sites *F* that will become free at the end of the generation is drawn from a binomial distribution with parameters *N* and *θ*. Individuals then reproduce in proportion to their fitness. That is, for each of the free sites in the population, we draw one mother individual and one father individual (which may incidentally be the same and then result in selfing) with weights proportional to their fitness. Mutations happen with probability *u* per generation per allele copy and then change an allele into the respective other allele. Finally, *F* adults are randomly killed to complete the generation.

### Mathematical analysis

Our first goal is to analytically get an understanding of the optimum number of 1 alleles (where 1 alleles correspond to extra metabolite, or upregulation of metabolites) for an individual and for the potential for genetic and thus chemical polymorphism in the population. To study the long-term fate of alleles and polymorphisms in a temporally variable environment with non-overlapping generations, geometric mean fitness over time is the appropriate fitness measure (Haldane and Jayakar 1963). It is given by

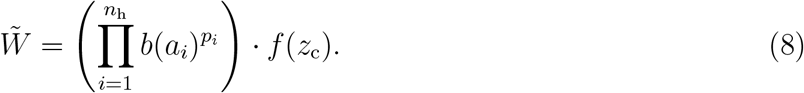

Note that this does not depend on the length *g* of phases, their ordering, or autocorrelation effects (Gillespie 1973). We use (8) to find the homozygous genotype (which has either 00 or 11 at each locus) with the highest fitness.

Next, we investigate whether and when there is scope for genetic polymorphism and thus also for variation in the chemical composition between individuals. For this, we use an invasion analysis similar to that in Wittmann *et al*. (2017). Given a fully monomorphic “resident” population where all individuals have the same multi-locus genotype and are homozygous at all loci, we determine whether or not a mutant with the respective other allele at a certain locus can invade (that is, increase in frequency in the long run when it starts out at very low frequency). Since we assume random mating, almost all copies of the rare allele are in heterozygotes and the number of heterozygotes is *N ·* 2*x ·* (1 *− x*) *≈* 2*Nx*, where *x* is the frequency of the rare allele. The rare allele increases in frequency if the number of heterozygotes increases in frequency. If *x*_*t*_ is the allele frequency at time *t*, the expected number of heterozygotes in the next generation *t* + 1 is approximately

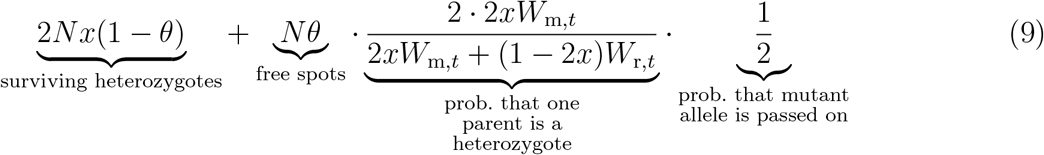

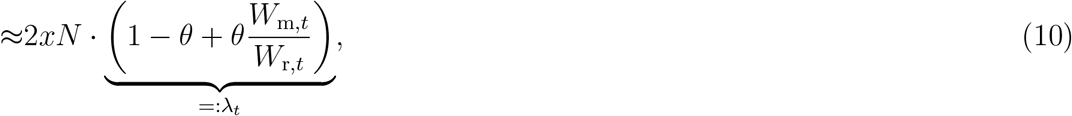

where *W*_m,*t*_ and *W*_r,*t*_ are the fitness of the mutant-resident heterozygote and of the resident homozygote at time *t*, as given by (7), and where we have assumed that 1 *− x ≈* 1 and that because *x* is close to zero, the denominator of the probability that a mutant is picked as a parent is approximately *W*_r,*t*_. Thus, the number of heterozygotes (and equivalently the frequency of the rare allele), grows by a factor *λ*_*t*_.

To check whether the mutant can invade, we then need to compute the geometric mean 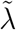 of *λ*_*t*_ over time. In the case of a single herbivore with occurrence probability *p*_1_, the mutant can invade if

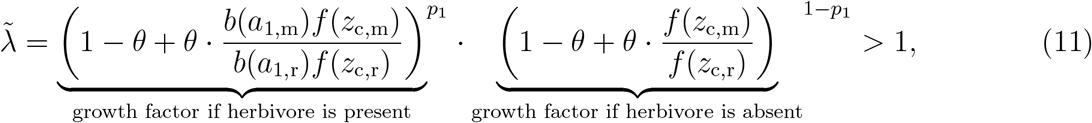

where the activity and cost traits are computed using (1), (2), (5), and (6). In the general case, we score polymorphism as possible if every homozygous resident type can be invaded by at least one mutant that differs from the resident at one locus (one could also consider mutants differing at multiple loci, but such mutants are very unlikely to arise for small mutation rates).

With non-overlapping generations (*θ* = 1) and the same effect size *m*_*l*1_ = 1 for all loci, (11) can be further simplified and there is scope for polymorphism if

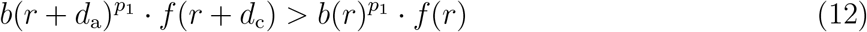

or

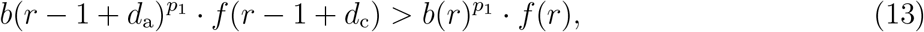

where *r* is the number of 11 loci in the resident. Note that in cases where *r* is 0 or *L*, only one of the possible mutants exists and so only one of the conditions needs to be checked.

The analysis can be generalised to the case of more than one herbivore:

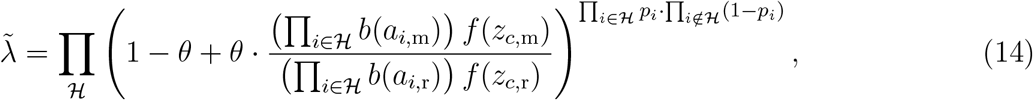

where the first product is over all possible subsets of herbivores *ℋ*. For example, with two herbivores

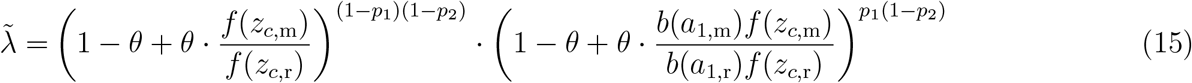

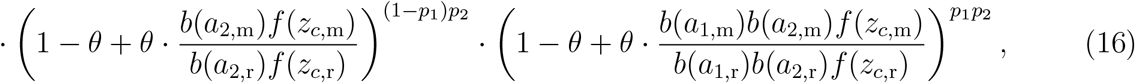

which we use below for the generalist-specialist-trade-off scenario.

### Individual-based simulations

The analytical results only inform us about the conditions under which the population would be expected to be fully monomorphic or not, i.e. somewhat polymorphic. But it cannot tell us at how many loci polymorphism will arise and how patterns of allele frequencies will look like also in the face of stochasticity, i.e. random genetic drift. Also, the analytical approach does not work with multiple herbivores and different effect sizes of the different loci against the different herbivores. Thus to investigate these aspects and characterise the resulting patterns of chemodiversity, for each parameter combination we ran 5 replicate individual-based simulations for 5000 generations with a population size *N* = 500, *g* = 5 generations per phase, a mutation rate of 0.0001, and 10 unlinked loci. Simulations were run for 5000 generations or 5500 generations (if there was an invasion, see below for a description of the invasion scenarios). The simulation model was implemented as a stochastic individual-based simulation in C++.

For each parameter combination we then quantified or estimated 1) the total number of “metabolites”, i.e. how many of the ten loci have some 1 alleles in the population (*γ* diversity), 2) the average number of “metabolites” per individual (also between 0 and 10), estimated as 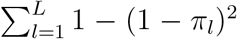, where *π*_*l*_ is the frequency of the 1 allele at locus *l* (*α* diversity), and 3) the average number of “metabolites” that are not shared in a randomly drawn pair of individuals (i.e. one individual has it, the other one does not), estimated as 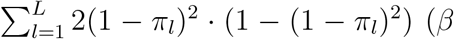 diversity). We evaluated each measure at the end of the simulation run and averaged across the 5 replicates.

## Results

### Testing the fluctuating selection and dominance reversal hypotheses

Let us first focus on the simplest case of a single herbivore (*n*_h_ = 1) with every locus having the same anti-herbivore effect *m*_*l*,1_ = 1. We will first address the two more general population genetic hypotheses, i.e., the fluctuating selection hypothesis and the dominance reversal hypothesis. For this, we tested whether and how dominance effects and fluctuating selection, two key factors for the maintenance of genetic diversity, affect chemodiversity patterns in our model (Fig. 2). To test for the effect of fluctuations, we compared three scenarios. In the fluctuating herbivory scenario (left column of Fig. 2), the herbivore was present in 20 % of phases and the baseline probability of escaping the herbivore when it is present was low (*b*_0_ = 0, blue curve in Fig. 1 a). In the constant high herbivory scenario (middle column), we used the same benefit function, but the herbivore was present in every phase, i.e., there were no fluctuations. In the constant low herbivory scenario (right column), the herbivore was also present in all phases, but the baseline probability to escape the herbivore was higher (green curve in Fig. 1 a) such that the overall herbivory pressure was similar to the fluctuating herbivory scenario. As might be expected because of the similar average herbivory pressure, in the fluctuating herbivory scenario and in the constant low herbivory scenario, the optimum number of metabolites based on geometric mean fitness was similar and lower than in the constant high herbivory scenario (upper row of Fig. 2).

**Figure 2.**
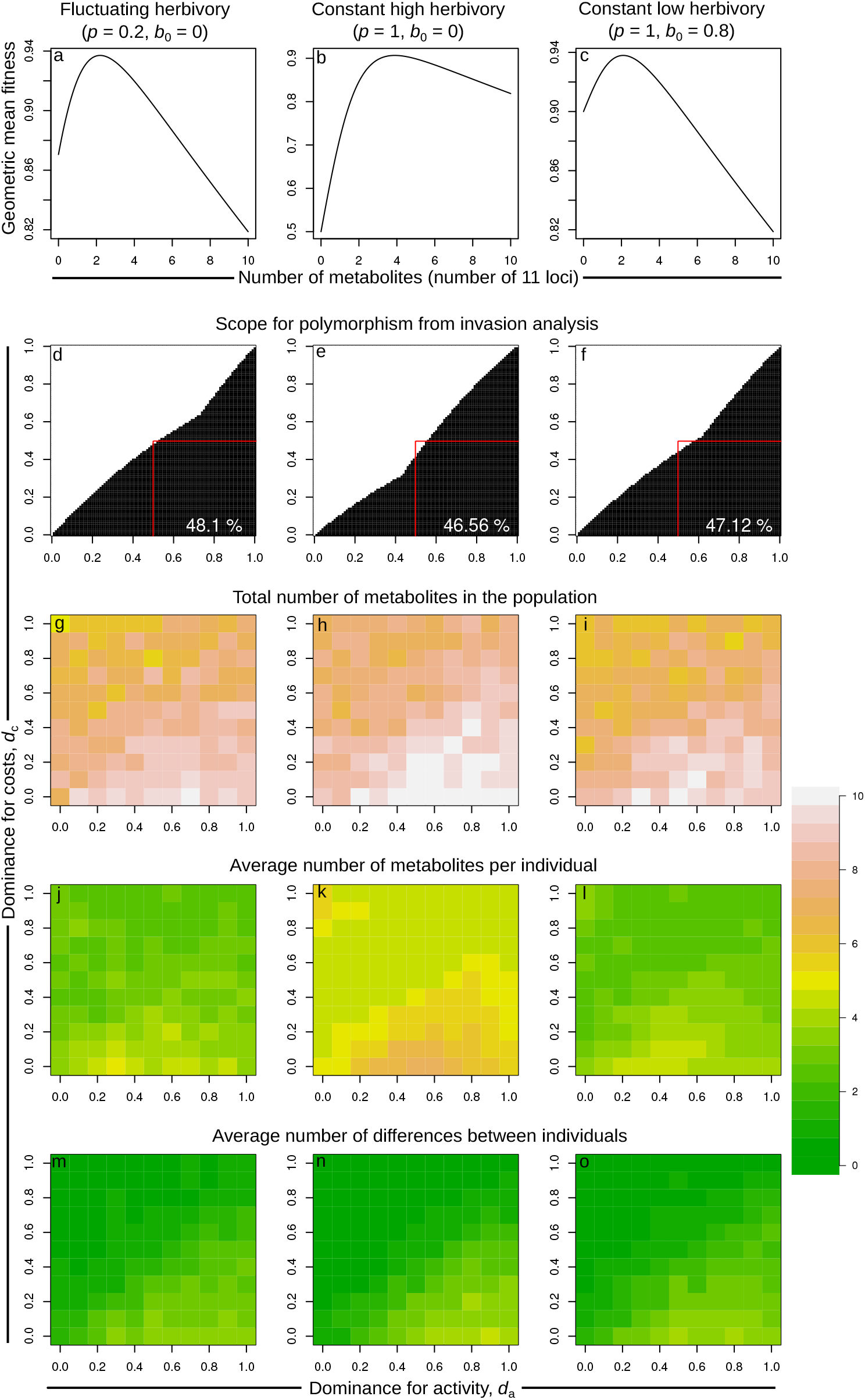
Dependence of chemodiversity patterns on dominance coefficients and fluctuations in the presence of one herbivore. a-c: Geometric mean fitness of homozygotes with different numbers of protection alleles. d-f: combinations of dominance coefficients for which the analytics predict that polymorphism is possible are shown in black. The proportion of dominance combinations for which polymorphism is possible is indicated as a percentage in the bottom right. The red square indicates the parameter space where polymorphism would be expected under the reversal of dominance hypothesis. g-o: Results of the corresponding stochastic individual-based simulations.

To test the dominance reversal hypothesis, in the second row of Fig. 2, we varied the dominance coefficients for benefits and costs independently. The black regions in this parameter space are regions where polymorphism is possible according to the analytic approach. The rough pattern is that polymorphism requires the dominance coefficient for activity *d*_a_ to be higher than the dominance coefficient for costs *d*_c_. Based on the dominance reversal hypothesis, one would have expected that polymorphism is only possible when the dominance for activity is larger than 0.5 and the dominance for costs is smaller than 0.5 (red squares). However, the actual coexistence region was larger, while also excluding some parameter combinations with *d*_a_ *>* 0.5 and *d*_c_ *<* 0.5. Against the expectation from the fluctuating selection hypothesis, the region where polymorphism is possible was only slightly higher in the fluctuating herbivory scenario compared to the two constant herbivory scenarios. That is, a sufficient difference between dominance for activity and dominance for costs can maintain polymorphism even in a constant environment.

The corresponding individual-based results (bottom three rows of Fig. 2) are qualitatively consistent with the analytical results. The total number of metabolites observed in the population (left column) and the average number of metabolites per individual (middle column) were higher in the constant high herbivory scenario compared to the other two scenarios. The average number of metabolites not shared between individuals (bottom row) was close to zero in regions where the analytical results predict that polymorphism is not possible, but reached values of four or more in regions where polymorphism is predicted to be possible. Also here, there were no large differences between the fluctuating herbivory and constant low herbivory scenario. The corresponding results for the case with overlapping generations where plants survive to the next time step with probability 0.7 (adult death probability *θ* = 0.3) are shown in Fig. S1. While the analytical results show no major differences, the simulation results indicate that with overlapping generations and adult mortality 0.3 per time step all three chemodiversity measures are slightly lower than with non-overlapping generations. However, if the adult mortality was further decreased (*θ* = 0.07) so that plants have an even longer average life span, all measures of chemodiversity are increased (Table S1). In both overlap scenarios, there was no clear difference in chemodiversity measures between the fluctuating herbivory scenario and the constant low herbivory scenario.

To test the hypothesis that a trade-off between generalist repulsion and specialist attraction, potentially with temporal fluctuations in specialist and generalist presence, promotes chemodiversity we ran a scenario with two herbivores where all ten metabolites had the same effect sizes and affected both the generalist herbivore (*m*_*l*1_ = 0.2, 0.5, or 1) and the specialist herbivore (*m*_*l*2_ = −0.2).

The results were qualitatively similar to those in Fig. 2 here and in other scenarios explored below. The difference between dominance for activity and dominance for costs seems to be the main driver for polymorphism. To better understand the effect of the generalist-specialist tradeoff, we summarized all the results by averaging the three quantities of interest (total metabolites, average number per individual, and average number of differences between individuals) across all dominance parameter combinations. We also obtained a standard error by first averaging for each replicate and then getting the standard error across replicates. Note that the standard errors are very small since although there are only five replicates per parameter combination, each replicate includes independent simulations for many dominance combinations such that stochastic effects mostly average out already within replicates. In addition we obtained the scope of polymorphism analytically as the % of dominance parameter space for which the analytical calculations predict that polymorphism is possible (cf. numbers in Fig. 2 d–f).

When the repellent effect on generalists was only as strong as the attractive effect on specialists (0.2), individuals produced few metabolites and there were few differences between individuals in metabolites produced (Fig. 3 b, c), consistent with no scope for polymorphism according to the analytic calculations (Fig. 3 d). For stronger repellent effects on generalist herbivores, the number of metabolites produced in the population and per individual as well as the number of differences between individuals increased. For high protective effects against generalists, the constant high herbivory scenario had the highest number of metabolites per individual, but the fewest differences between individuals according to both simulations and analytical results. Again, there were no consistent differences between the fluctuating herbivory and constant low herbivory scenarios. The analytical results (Fig. 3 d) suggest that the generalist-specialist trade-off can slightly widen the dominance region where polymorphism is possible (compare to Fig. 2). Since the scope for polymorphism was in some cases above 50 %, there are some scenarios where polymorphism is possible although dominance for costs is slightly higher than dominance for activity.

**Figure 3.**
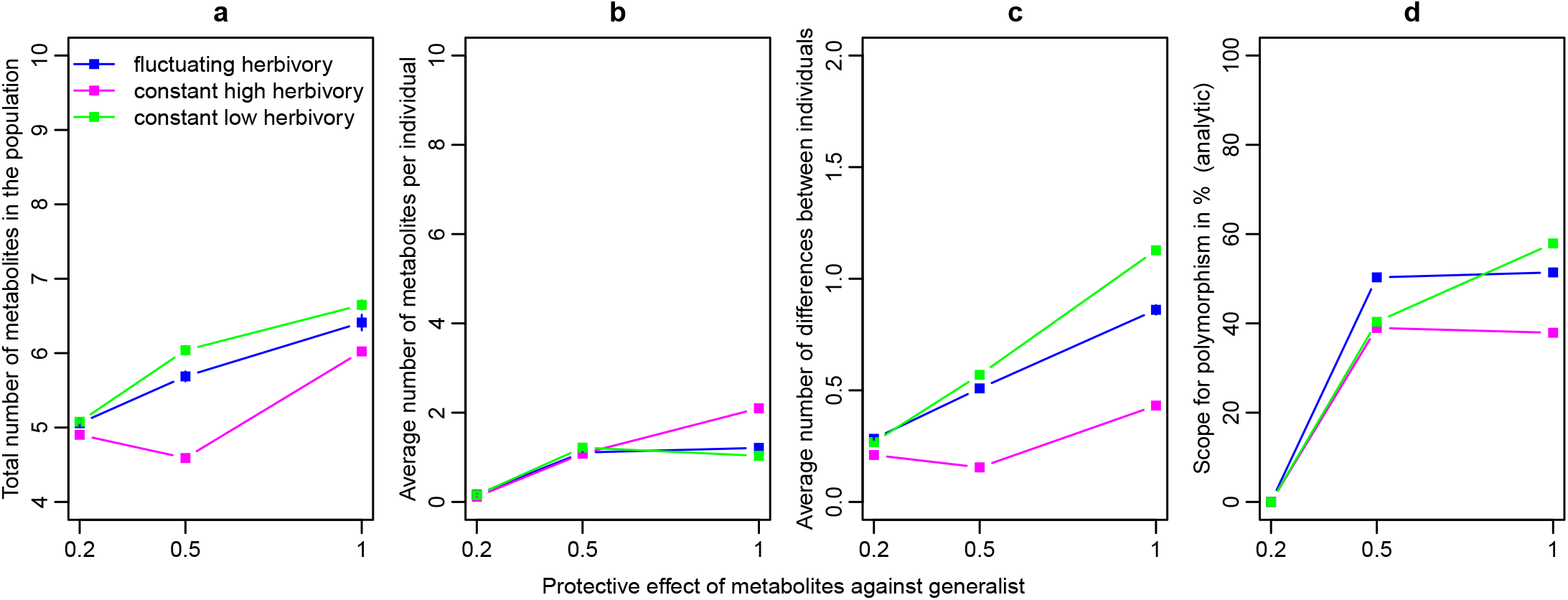
Patterns of chemodiversity in a scenario with one specialist which is attracted by metabolites with effect size -0.2 and one generalist. The magnitude of the protective effect against the generalist is varied on the x axis. The bars indicate the standard error, but many of them are so small that they fall entirely inside the point and are therefore not visible. Note that the points are connected for increased visual clarity, but that nonlinear and even nonmonotonic relationships are possible.

Curiously, in the constant high herbivory scenario, the total number of metabolites in the population and the average number of differences between individuals was smallest at intermediate protective effect against generalists. A closer look at the underlying allele frequency distributions (Fig. S2 b) shows that in this scenario, there was usually one locus where the presence allele was fixed or at very high frequency, whereas most other loci were at 0 or very low frequency. For lower or higher protective effect against generalist herbivores, there were more loci with allele frequencies between 0.01 and 0.1 that thus contributed more to the total number of loci in the population and to individual differences. From these results, it is not clear why the allele frequency distribution in this case was so different, but it could be that in this scenario genotypes with two presence alleles at one locus are close to optimal, so that other genotypes are more strongly selected against compared to other scenarios.

### Testing the interaction diversity hypothesis

Next, to explore patterns of chemodiversity predicted under the interaction diversity hypothesis, we ran simulations with one, two or five herbivores. In addition to the three fluctuation (or not) scenarios, we considered two scenarios for the effect size distributions: In the “only repellent” scenario, each locus had protective effects 0.2, 0.4, 0.6, 0.8 or 1 against each herbivore with probability 0.1 each, and was neutral with probability 0.5. In the “repellent + attractive” scenario, the probabilities for the different protective effect sizes were the same, but each metabolite also had attractive effects of magnitude 0.2, 0.4, 0.6, 0.8, 1 with probability 0.02 each, and the probability to be neutral was then only 0.4 (see Fig. S3 for a visualisation of the two distributions). The values were independently drawn from these distribution for each combination of locus, herbivore, and replicate. That is, a single metabolite could have different effects on the different herbivores.

As predicted under the interaction diversity hypothesis, the average number of metabolites per individual increased with the number of herbivores, albeit only very weakly under constant low herbivory (Fig. 4 b). Although not a straightforward prediction of the interaction diversity hypothesis, our model allows us to also predict the effect of number of herbivores on the other two aspects of chemodiversity. The average number of differences between individuals increased with number of herbivores under all scenarios. The effect of number of herbivores on the total number of metabolites in the population differed among scenarios: more herbivores led to more metabolites under constant high herbivory, as expected under the interaction diversity hypothesis. Under fluctuating herbivory, the number of metabolites in the population stayed roughly constant with increasing numbers of herbivores. Unexpectedly, an increasing number of herbivores slightly decreased metabolite numbers under constant low herbivory in the repell + attract scenario. One contributing factor to this appears to be that with increasing herbivore numbers the probability for a metabolite to attract at least one herbivore increases (0.1 for 1 herbivore, 0.19 for 2 herbivores, and 0.41 for 5 herbivores), while the probability that such loci are lost stays roughly constant (0.49, 0.48, 0.52). Thus, with more herbivores, there are fewer metabolites that are unconditionally beneficial. This can lead to a higher probability of presence alleles to be lost in the population and therefore lower total numbers of metabolites in the population.

**Figure 4.**
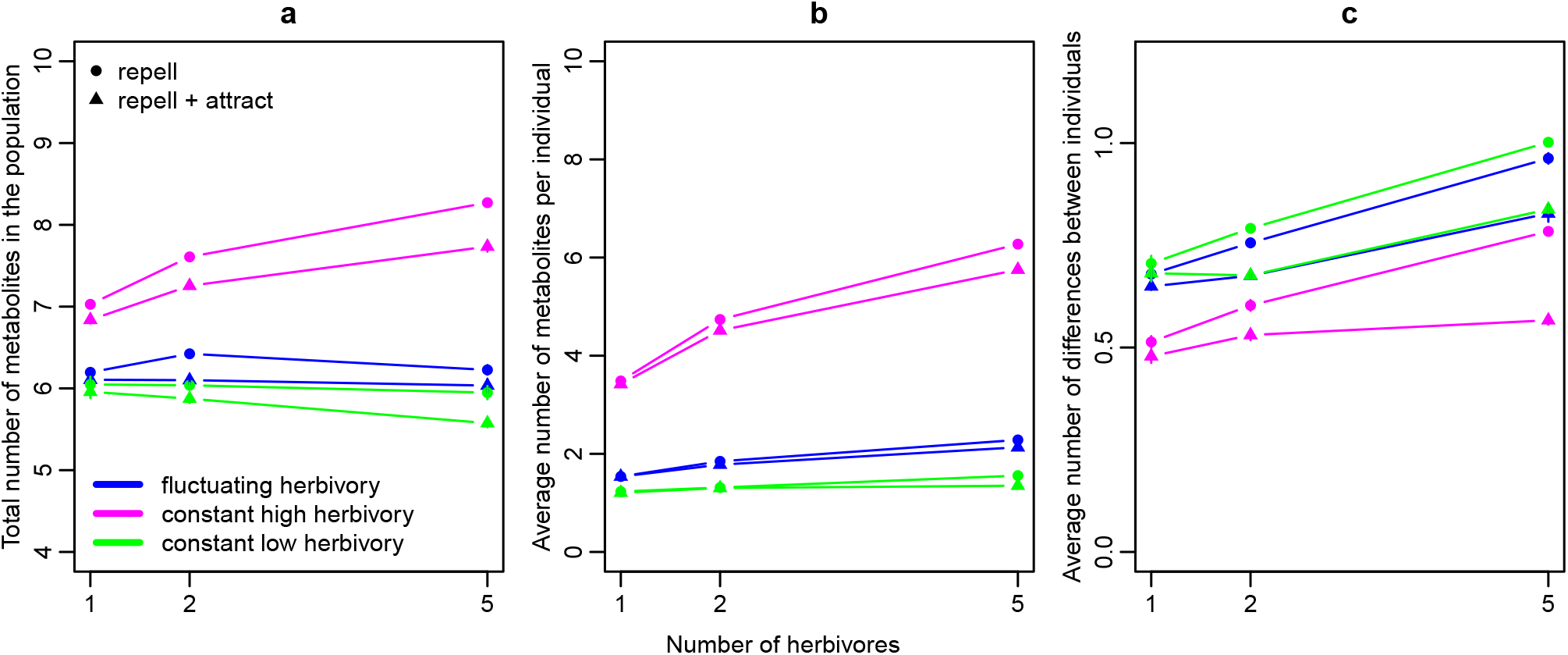
Effect of the number of herbivores and the distribution of effect sizes on chemodiversity patterns. The bars indicate the standard error, but many of them are so small that they fall entirely inside the point and are therefore not visible. Note that the points are connected for increased visual clarity, but that nonlinear and even nonmonotonic relationships are possible.

### Testing the synergy hypothesis

To explore patterns of chemodiversity under the synergy hypothesis, we focused on a single herbivore and compared different shapes of the benefit function (small plots below Fig. 5). Scenarios with a low half-saturation constant *a*_half_ had diminishing returns of anti-herbivore protection with increasing number of metabolites, whereas scenarios with high half-saturation constant had a synergistic benefit function such that over most of the range, the benefit of having multiple metabolites was larger than the benefit expected from single metabolite effects. Had we chosen *b*_0_ = 0, like in most other simulations, the scenarios would have differed not only in the shape but more strongly also in the overall protection level against herbivores. Thus, the parameter *b*_0_ was chosen for each scenario such that individuals with activity 0 had the same probability 0.01 to escape the herbivore when it is present.

**Figure 5.**
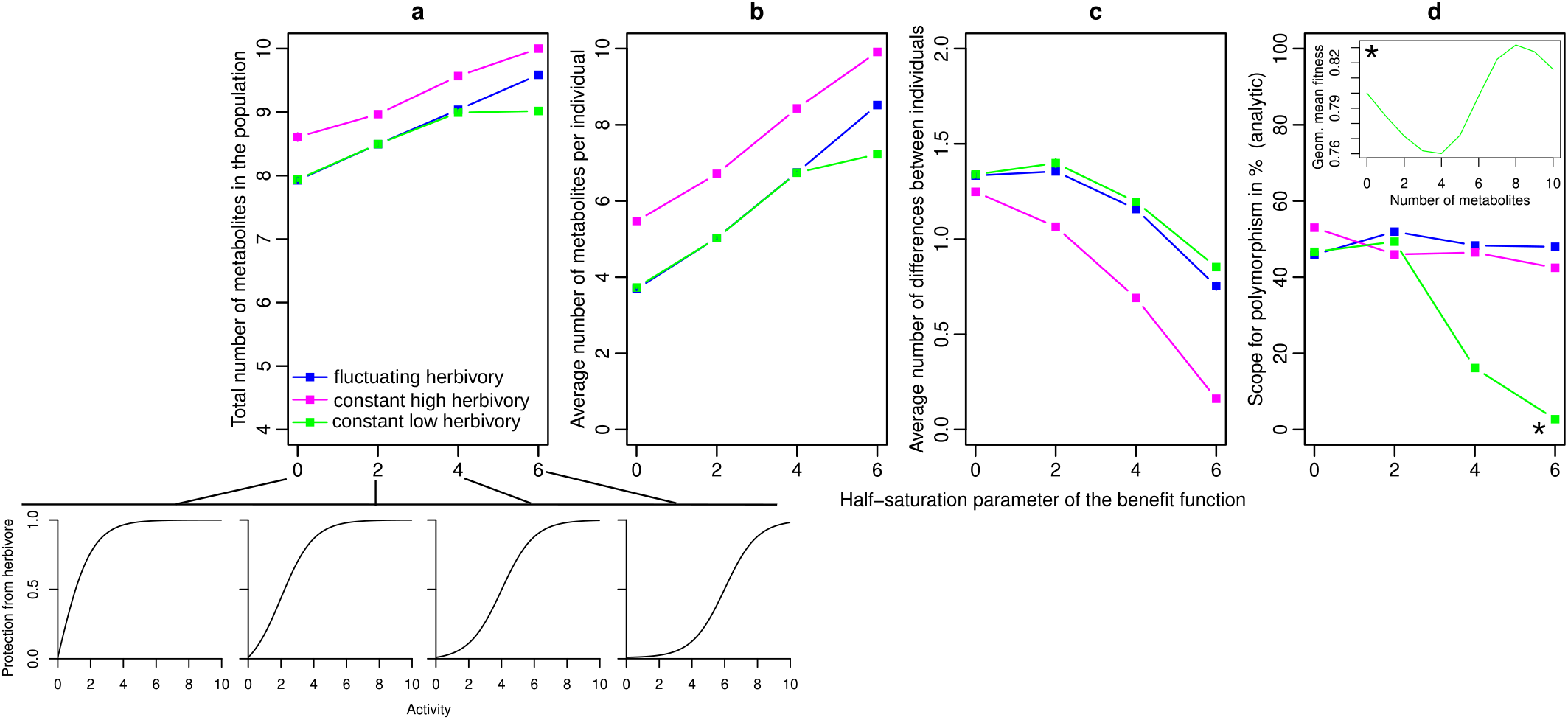
Effect of the shape of the benefit function on patterns of chemodiversity in the case of a single herbivore. With increasing half-saturation parameter, the benefit function becomes more synergistic (see small plots below a). The bars indicate the standard error, but many of them are so small that they fall entirely inside the point and are therefore not visible. The inset in (d) shows geometric mean fitness for the various homozygote genotypes with different number of metabolites for the constant-low herbivory scenario with *a*_half_ = 6 (see ^*^). Note that the points are connected for increased visual clarity, but that nonlinear and even nonmonotonic relationships are possible.

We observe that both under constant or fluctuating herbivory, the total number of metabolites in the population and the average number of metabolites per individual increase with the half-saturation parameter of the benefit function, i.e., as the benefit function becomes more synergistic. In contrast, the average number of differences between individuals decreases as the benefit function becomes more synergistic. The fluctuating herbivory and constant low herbivory scenarios produced almost the same chemodiversity patterns, but interestingly they diverged for very strongly synergistic benefit functions where the total number and average number per individual were higher under fluctuating herbivory. The constant high herbivory scenario had the highest number of metabolites, both in total and per individual, but the lowest number of differences between individuals. The analytic approximation (Fig. 5 d) did not capture the between-individual chemodiversity very well and predicted that the scope for polymorphism is lowest in the constant low herbivory scenario. The reason seems to be that the analytical invasion approach only checks whether each homozygous genotype can be invaded by one of the other genotypes that differs at only one position. This corresponds to a very small mutation rate. For the synergistic benefit function, the geometric mean fitness function can be non-monotonic (see inset), suggesting that the extreme genotype that does not produce any metabolites might be non-invasible by all one-step mutants, but could be invasible by mutants differing in multiple positions. In the simulations, however, the mutation rate seems to be large enough that multiple mutants differing in several positions segregate in the population. This could allow other genotypes to invade the no-metabolite genotype in many cases where the analytical results suggest it is not possible.

### Testing the screening hypothesis

Because our model does not explicitly include metabolic pathways, most of the predictions of the screening hypothesis cannot be tested and require a more complex model. However, there is one prediction we can test. Based on the screening hypothesis, plants with a higher number of metabolites should have a higher probability of having a metabolite that is active against a newly colonizing herbivore and therefore a higher fitness after such an invasion (Jones and Firn 1991). To test this, we ran simulations under the fluctuating herbivory scenario and with varying dominance parameters, where 1, 2, or 5 herbivores were present until time 5000 and an additional herbivore was introduced at time 5000 and present until the end of the simulation at time 5500. For these simulations, we used the distributions of effect sizes from Fig. S3 and the effect of each metabolite on each herbivore was drawn independently, also for the additional herbivore.

Confirming the expectation from the screening hypothesis, we found that plant populations with a higher chemodiversity (independently of whether it was measured as total metabolites produced, average number of metabolites produced per individual, or average number of differences between individuals), maintained higher average fitness in the time period after the invasion of an additional herbivore (Fig. 6). This was the case both when metabolites had only repellent effects (top row of Fig. 6) and when metabolites had both repellent and attractive effects (bottom row of Fig. 6).

**Figure 6.**
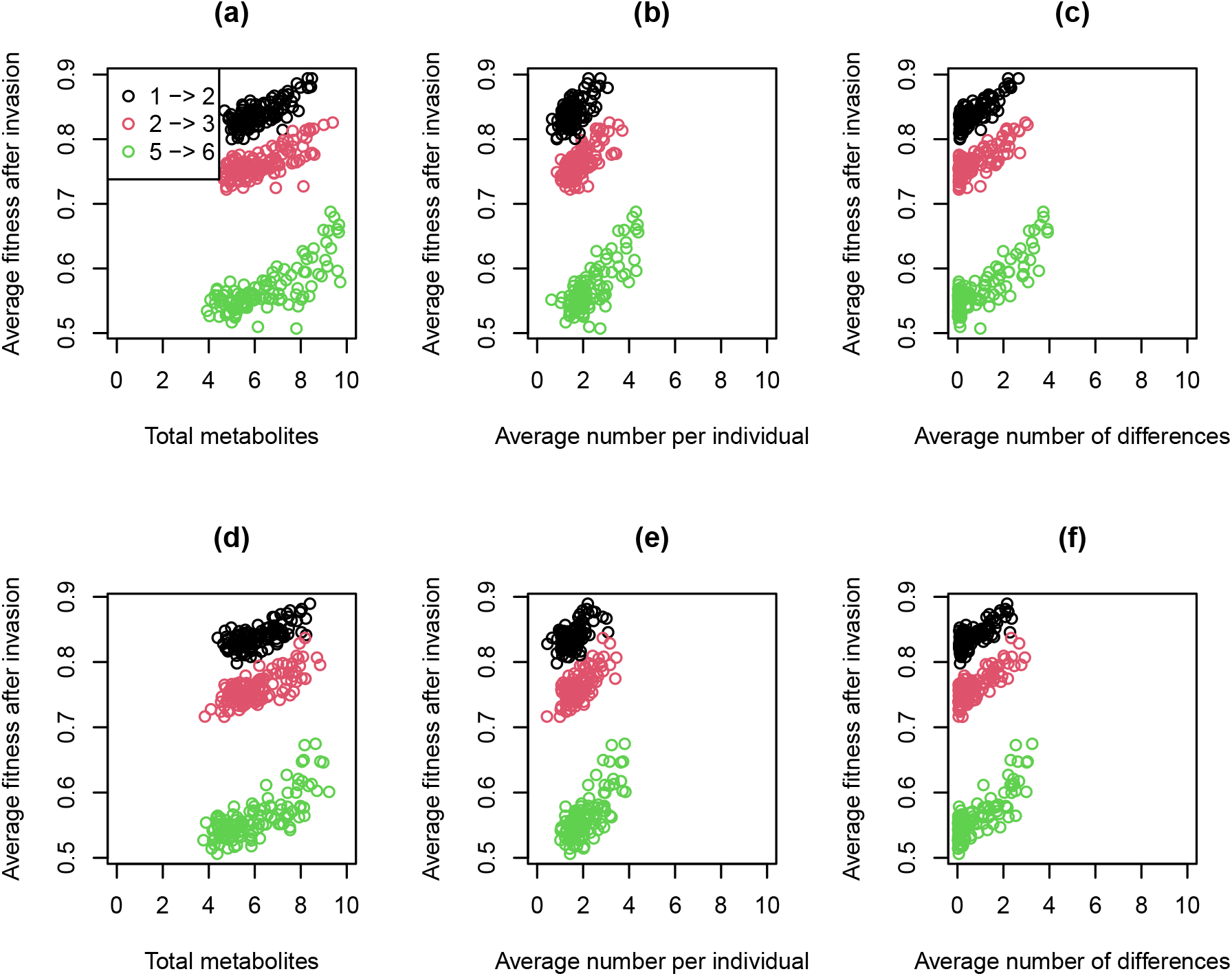
Average fitness over 500 generations after the invasion of a new herbivore as a function of the different aspects of chemodiversity before invasion (average over the last 95 generations before invasion). Each point corresponds to the average over five replicates for one of the dominance and herbivore number scenarios. Top row: Scenario with only repellent effects of metabolites on herbivores as depicted in Fig. S3 a. Bottom row: Scenario with both repellent and attractive effects of metabolites on herbivores as depicted in Fig. S3 b.

## Discussion

### Support for and new predictions of the five hypotheses

We have presented a stochastic population genetics modeling approach that provides a quantitative testing ground for various verbal models and hypotheses on the evolution of chemodiversity.

Our analytical and simulation results agree with the dominance reversal hypothesis in that differences in dominance between different traits, here the costs and benefits of defense metabolites, are a key factor for the maintenance of polymorphism. However, the exact conditions for maintenance of polymorphism at defense loci did not match with the hypothesis. We had expected polymorphism when dominance for costs is below 0.5 and dominance for activity is above 0.5. But polymorphism actually emerged when the dominance for anti-herbivore activity was substantially larger than the genetic dominance for the costs of producing metabolites, i.e. heterozygotes experienced relatively large benefits and relatively low costs of their one copy of the metabolite-producing allele. This led to a high total number of metabolites produced in the population and a high average number of differences between individuals. A reversal of dominance was not necessary, but also not completely sufficient for the maintenance of polymorphism.

Antagonistic pleiotropy with changes in dominance between traits has previously shown to be a powerful mechanism for the maintenance of genetic polymorphism (Curtsinger *et al*. 1994; Rose 1982). Although the focus has been mostly on reversals of dominance, a closer examination of the model by Rose (1982) shows that also in a simpler model with antagonistic pleiotropy at a single locus in a constant environment, smaller quantitative differences in dominance between traits can be sufficient to maintain polymorphism (Supporting Information Section 1). Similar observations were made by Van Dooren (2006) and Brud (2023). Thus, one should talk less about beneficial dominance reversals and more about beneficial dominance shifts as a factor promoting genetic variation.

Differences in dominance among affected traits for pleiotropic loci have been observed for example for loci influencing mandibular morphology in mice (Ehrich *et al*. 2003) and can arise in branched enzyme pathways with nonlinearities and feedbacks (Keightley and Kacser 1987). In fact, since many traits and fitness components emerge from highly nonlinear processes, there is no particular reason why dominance should be equal for all traits affected by a pleiotropic locus. For metabolic pathways in particular, heterozygotes might be expected to be intermediate with respect to the costs of genes and enzymes (dominance 0.5). Flux through the respective pathway is often a concave (diminishing returns) function of enzyme concentration (Kacser and Burns 1981). Thus heterozygotes will be closer in flux to the homozygote with the larger flux and dominance for activity is expected to be larger than 0.5 (Kacser and Burns 1981). This is the case as long as the enzyme is not saturated with substrate (Cornish-Bowden 1987; Kacser and Burns 1981; Wright 1934). This appears realistic since in specialized metabolism, flux through a pathway is frequently controlled at gateway entry points such as phenylalanine ammonia lyase for phenylpropanoids, chalcone synthase for flavonoids and 3-hydroxymethylglutaryl coenzyme A reductase for terpenoid biosynthesis (Dixon and Dickinson 2024) rather than enzymes deeper in the pathway. Also, compared to enzymes in primary metabolism, enzymes in specialised metabolism have on average less optimized kinetic parameters, suggesting that selection pressure on efficiency in specialized metabolism is relatively low, presumably because of the lower flower through these pathways (Bar-Even *et al*. 2011; Bar-Even and Salah Tawfik 2013). Thus it appears plausible that enzymes have not evolved Michaelis constants *K*_*M*_ low enough to match the physiological substrate concentrations and thus do not operate at saturation, making dominance for activity likely. Although changes and in particular reversals in dominance have been discussed for other types of trait in evolutionary biology, to our knowledge, this is the first time that dominance changes are highlighted as a potential key factor in the evolution of plant chemodiversity.

Contrary to the expectation from the fluctuating selection hypothesis, temporal fluctuations in herbivore presence had surprisingly small effects for chemodiversity in our model. The reason seems to be that selection pressures under fluctuating herbivore occurrence are in the long run quite similar to scenarios with continuous presence of herbivores but a lower herbivory pressure (Figs. 2 and 4). If previous studies on the interplay of temporally fluctuating selection and dominance changes (Hedrick 1976; Wittmann *et al*. 2017) had also included such a constant control treatment, we speculate that they might have found similar maintenance of polymorphism without fluctuations. However with multiple herbivores or strong synergy, fluctuating herbivore occurrence slightly promoted chemodiversity compared to constant low herbivory both within individuals and within populations, but not between individuals (see Figs. 4 and 5). In a model with antagonistic pleiotropy driven by a fecundity-viability trade-off of flowering time, Brown and Kelly (2018) similarly found only a small effect of environmental fluctuations on the maintenance of polymorphism. Previous theory has shown that temporally fluctuating selection can more easily maintain genetic variation if generations are overlapping (Chesson and Warner 1981). Thus we had hypothesized that generation overlap would increase the parameter space where temporal fluctuations promote chemodiversity, but at least for the scenarios we investigated, this does not appear to be the case. For overlapping generations with a relatively large adult death probability, all chemodiversity measures were reduced while with a smaller adult death probability and thus more generation overlap, all chemodiversity measures were increased (Table S1). Thus, we would predict long-lived perennials to harbor more chemodiversity than annuals. However, long generation overlap increased chemodiversity in all herbivory scenarios with or without fluctuations. Thus, generation overlap did not increase the power of fluctuating selection to maintain chemodiversity. However, we have assumed here that herbivory only influences plant reproduction, but not the probability to survive to the next time step, which may not be realistic. More detailed modeling is needed to clarify how plant life history and fluctuating herbivory interact to shape chemodiversity.

As expected from the interaction diversity hypothesis (Whitehead *et al*. 2021), with increasing number of herbivores, the average number of metabolites per individual increased (see Fig. 4). Surprisingly, this did not always lead to a higher total number of metabolites in the population, apparently because some of the scenarios with more herbivores also caused a larger probability of loss of rare alleles. Maybe the most interesting result of the simulations with multiple herbivores is that increasing herbivore numbers do not just promote an increase in the number of metabolites per individual, a straightforward prediction, but also lead to more differences in metabolite composition between individuals, a non-trivial prediction that would not be possible without quantitative models such as ours.

Consistent with the synergy hypothesis, our model produced generally larger numbers of metabolites in the population and per individual when metabolites had synergistic effects on protection against herbivores (Fig. 5). However, the average number of differences between individuals decreased with increasing synergy. This is also expected since when most individuals have most metabolites, they cannot differ in many metabolites.

Lastly, our simulations results support one of the predictions of the screening hypothesis, namely that plant populations with higher chemodiversity before invasion cope better with the invasion of an additional herbivore with randomly drawn traits. Note however, that this benefit of chemodiversity is not the reason for the evolution of chemodiversity in our model. Chemodiversity evolved in response to the “native” herbivores and if, because of the parameter settings, a higher diversity evolved, this coincidentally also helped to cope with a new herbivore.

In summary, almost all hypotheses yielded new unexpected predictions for one or more aspects of chemodiversity. For instance, we have found that although both increasing the number of herbivores and increasing the degree of synergy leads to increases in the number of metabolites per individual, at least for the parameter combinations we considered a higher number of herbivores lead to more differences in metabolites between individuals, whereas increasing synergy leads to fewer differences between individuals.

### Limitations and outlook

Here we have taken a simplified approach where the genotype directly determines whether or not a metabolite is produced. However, in reality the link between genotype and chemotype may not be so direct. For example, if metabolites are produced by multi-step enzymatic pathways, a metabolite can only be produced if the enzyme catalyzing its synthesis from its direct precursor is expressed, but also all precursors need to be present and the enzymes necessary to produce them need to be expressed. Thus metabolites produced in complex pathways do not evolve independently. A logical next step is to develop more mechanistic models that do not take the shortcut from genotype to chemotype, but model the underlying proteome and metabolic pathways. Such models could then be used to test other predictions of the screening hypothesis and to model quantitative variation in addition to qualitative presence-absence variation.

Second, we have not taken into account here that plants may be affected by the chemical profile and repellent or attractive effects of their neighbors, either through associational resistance or associational susceptibility. These phenomena can under some conditions create negative fre-quency dependent selection and then also contribute to maintaining chemodiversity in the sense of differences between individuals (Sato 2018).

Finally, here we have assumed for simplicity that plant population sizes are constant and overall herbivore pressures are independent of evolution in the plant population. It would also be interesting to include population dynamics of plants and herbivores as well as plant-herbivore coevolution. Eco-evolutionary feedbacks e.g. with specialist herbivores may be very important for the evolution of plant chemodiversity and should be addressed in future models.

## Acknowledgements

We thank Anke Steppuhn and other members of the research unit FOR 3000 for helpful discussions and Frans Thon for constructive comments on the manuscript.

## Author contributions

MJW and AB designed the research. MJW developed, programmed and analysed the model, and wrote the manuscript, with input from AB.

## Data availability

Simulation code and analysis scripts are provided in a supplementary zip folder.

## Supporting Information legends

**Table S1** Average chemodiversity levels across all dominance parameter combinations for different herbivory scenarios and with different adult death probabilities *θ*.

**Figure S1** Dependence of chemodiversity patterns on dominance coefficients and fluctuations in the presence of one herbivore and with overlapping generations with *θ* = 0.3.

**Figure S2** Allele-frequency spectra for generalist-specialist scenarios and different fluctuation scenarios.

**Figure S3** Effect size distributions used for simulations with multiple herbivores.

**Supporting Information Section 1** Conditions for maintenance of polymorphism at a single locus with antagonistic pleiotropy in a constant environment

**Table S2** Fitness effects in a simple single-locus antagonistic pleiotropy model with two fitness components (*W*_1_ and *W*_2_) that act either additively or multiplicatively.

**Figure S4** Changes in dominance can maintain polymorphism at a single locus with antagonistic pleiotropy, even without reversal of dominance.

**Code S1** Zip folder containing simulation code and analysis scripts.

## Supporting information

**Table S1.** Average chemodiversity levels across all dominance parameter combinations for different herbivory scenarios and with different adult death probabilities *θ*. TMP = average total number of metabolites in the population, MPI = average number of metabolites per individual, DM = average number of differences between individuals.

| Herbivory scenario | $\theta$ | TMP | MPI | DM |
| --- | --- | --- | --- | --- |
| Fluctuating | 1 | 7.59 | 2.91 | 1.31 |
| Constant high | 1 | 8.27 | 4.70 | 1.26 |
| Constant low | 1 | 7.50 | 2.87 | 1.35 |
| Fluctuating | 0.3 | 6.55 | 2.87 | 1.01 |
| Constant high | 0.3 | 7.41 | 4.59 | 1.01 |
| Constant low | 0.3 | 6.46 | 2.77 | 1.04 |
| Fluctuating | 0.07 | 7.74 | 3.64 | 2.02 |
| Constant high | 0.07 | 8.74 | 5.48 | 2.02 |
| Constant low | 0.07 | 7.72 | 3.56 | 2.00 |

**Figure S1.**
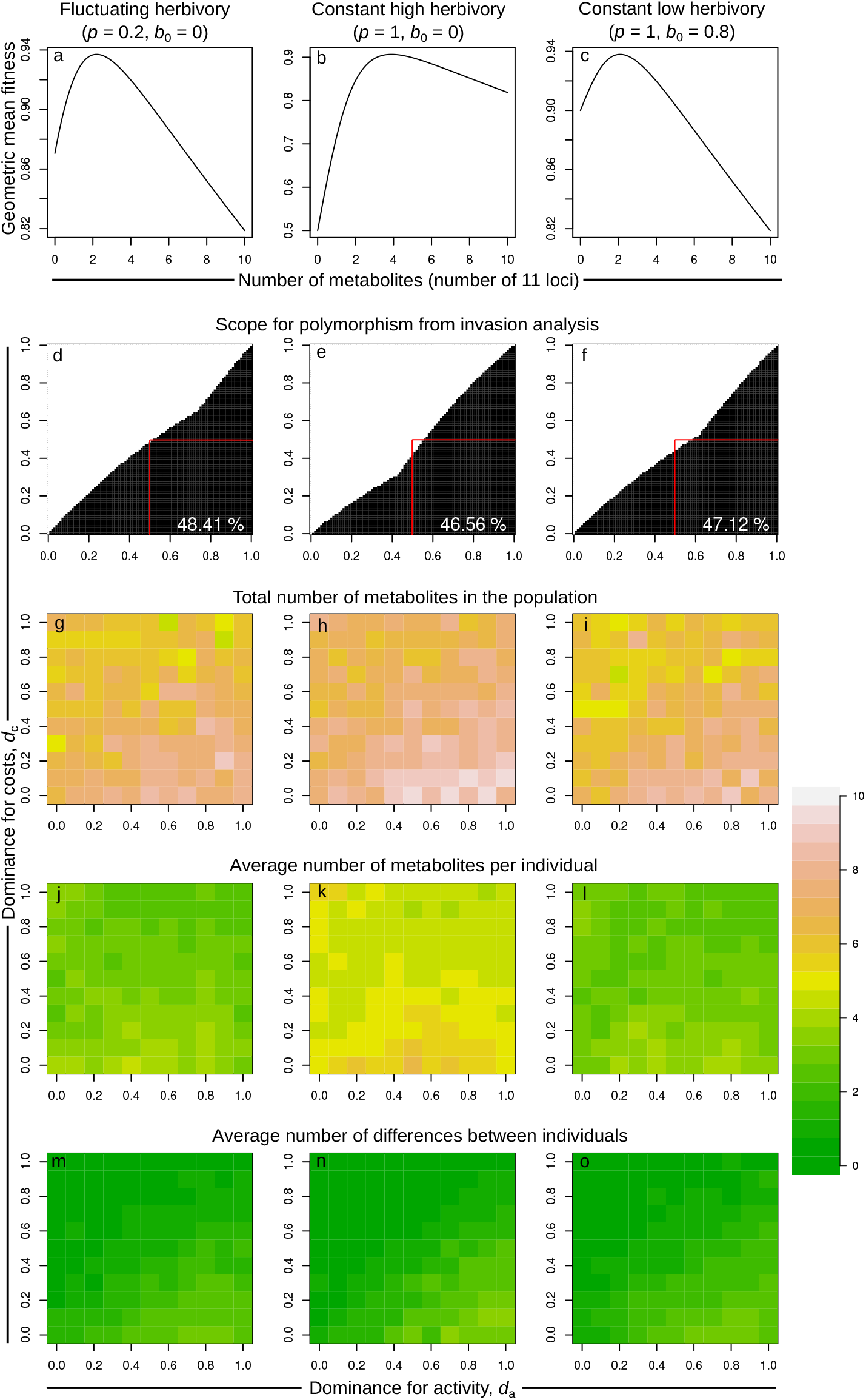
Dependence of chemodiversity patterns on dominance coefficients and fluctuations in the presence of one herbivore and with overlapping generations with *θ* = 0.3. a-c: Geometric mean fitness of homozygotes with different numbers of protection alleles. d-f: combinations of dominance coefficients for which the analytics predict that polymorphism is possible. The proportion of dominance combinations for which polymorphism is possible is indicated as a percentage in the bottom right. g-o: Results of the corresponding stochastic individual-based simulations.

**Figure S2.**
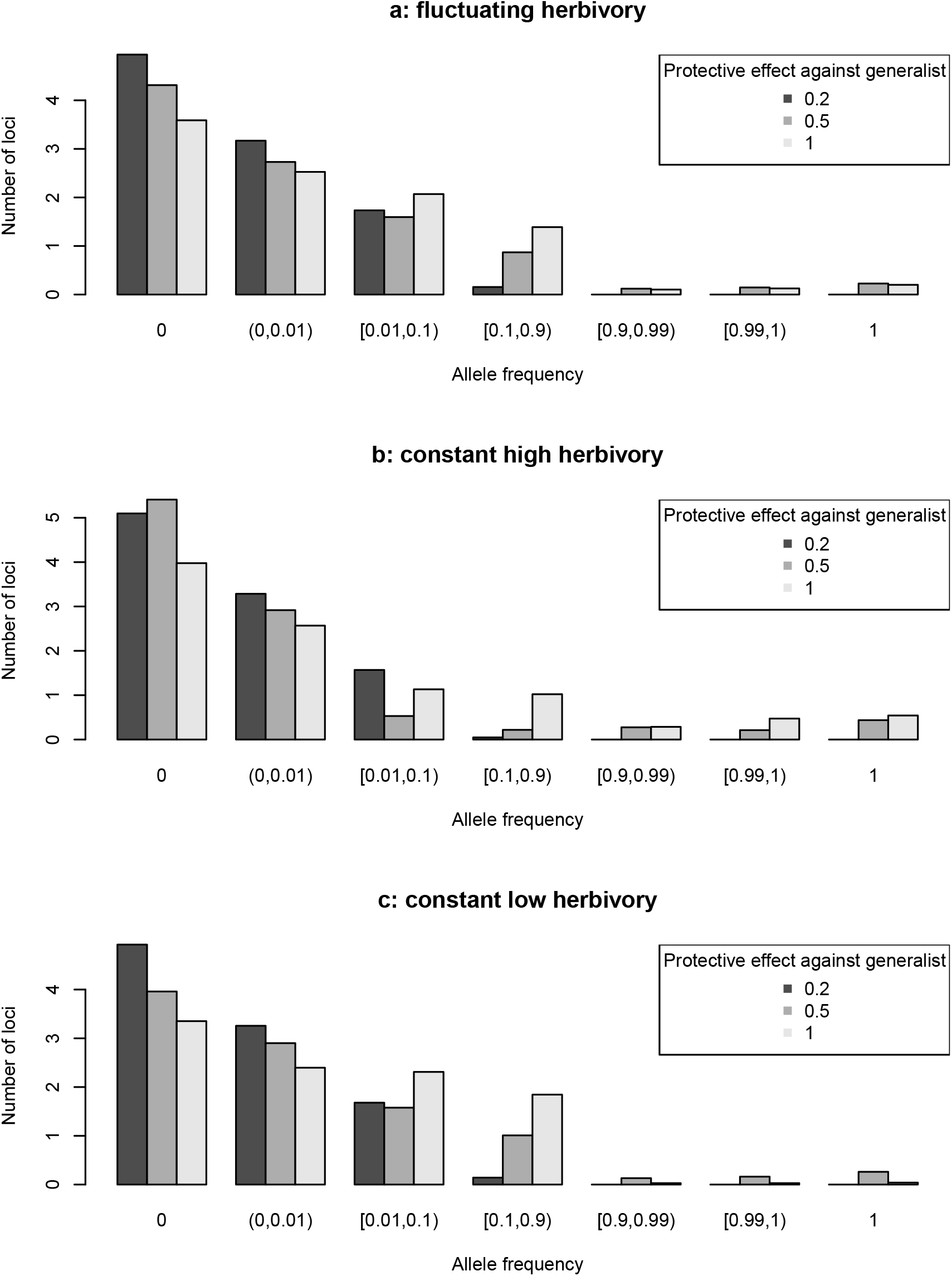
Allele-frequency spectra for generalist-specialist scenarios and different fluctuation scenarios. The underlying simulation results are the same as in Fig. 3. Each bar represents the number of loci (out of 10) that have allele frequencies in a given interval at the end of the simulation, averaged over all dominance parameter combinations and over the five replicates.

**Figure S3.**
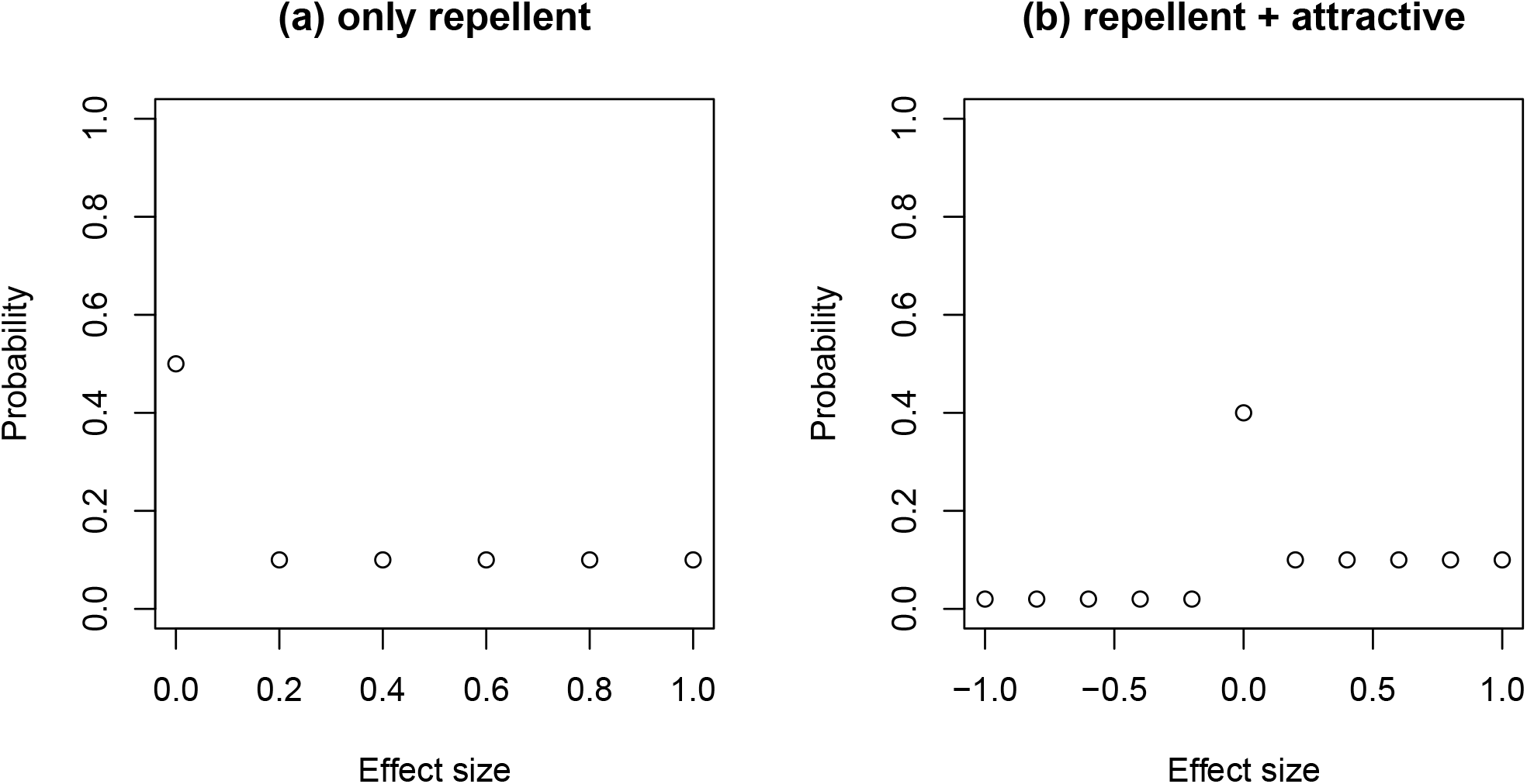
Effect size distributions used for simulations with multiple herbivores. Positive values correspond to protective effects against herbivores, whereas negative values correspond to attractive effects.

### Supporting Information Section 1

#### Conditions for maintenance of polymorphism at a single locus with antagonistic pleiotropy in a constant environment

Here we examine the single-locus antagonistic pleiotropy model by Rose (1982), but parameterized slightly differently so that it becomes more comparable to our model (Table S2). There are two alleles at the locus, 0 and 1. Allele 1 increases fitness component 1, *W*_1_, but decreases fitness component 2, *W*_2_. Like Rose (1982), let us consider the cases of additive selection and multiplicative selection. In each case, polymorphism is maintained if the total fitness (*W*_1_ + *W*_2_ in the additive case or *W*_1_*W*_2_ in the multiplicative case) of heterozygotes is higher than that of either homozygote. The dominance combinations for which this is the case are shown in Fig. S4. The regions that allow for polymorphism look qualitatively similar to those in our model. As in our model, there are combinations where polymorphism is possible even though there is no reversal of dominance. It is sufficient if the dominance for the benefits of the 1 allele on *W*_1_ is substantially larger than the dominance for the negative effect of the 1 allele on *W*_2_. The parameter region with polymorphism is largest if selection strength is equal on both fitness components (middle column in Fig. S4). If selection is stronger on fitness component 1, the polymorphism region is reduced to combinations where dominance for fitness component 1 is high. And when selection is stronger on fitness component 2, it is is reduced to combinations where dominance for fitness component 2 is low.

**Table S2.** Fitness effects in a simple single-locus antagonistic pleiotropy model with two fitness components (*W*_1_ and *W*_2_) that act either additively or multiplicatively.

| Genotype | 11 | 10 | 00 |
| --- | --- | --- | --- |
| $W_1$ | $1 + s_1$ | $1 + d_1s_1$ | 1 |
| $W_2$ | $1 - s_2$ | $1 - d_2s_2$ | 1 |
| $W_1 + W_2$ | $2 + s_1 - s_2$ | $2 + d_1s_1 - d_2s_2$ | 2 |
| $W_1W_2$ | $(1 + s_1)(1 - s_2)$ | $(1 + d_1s_1)(1 - d_2s_2)$ | 1 |

**Figure S4.**
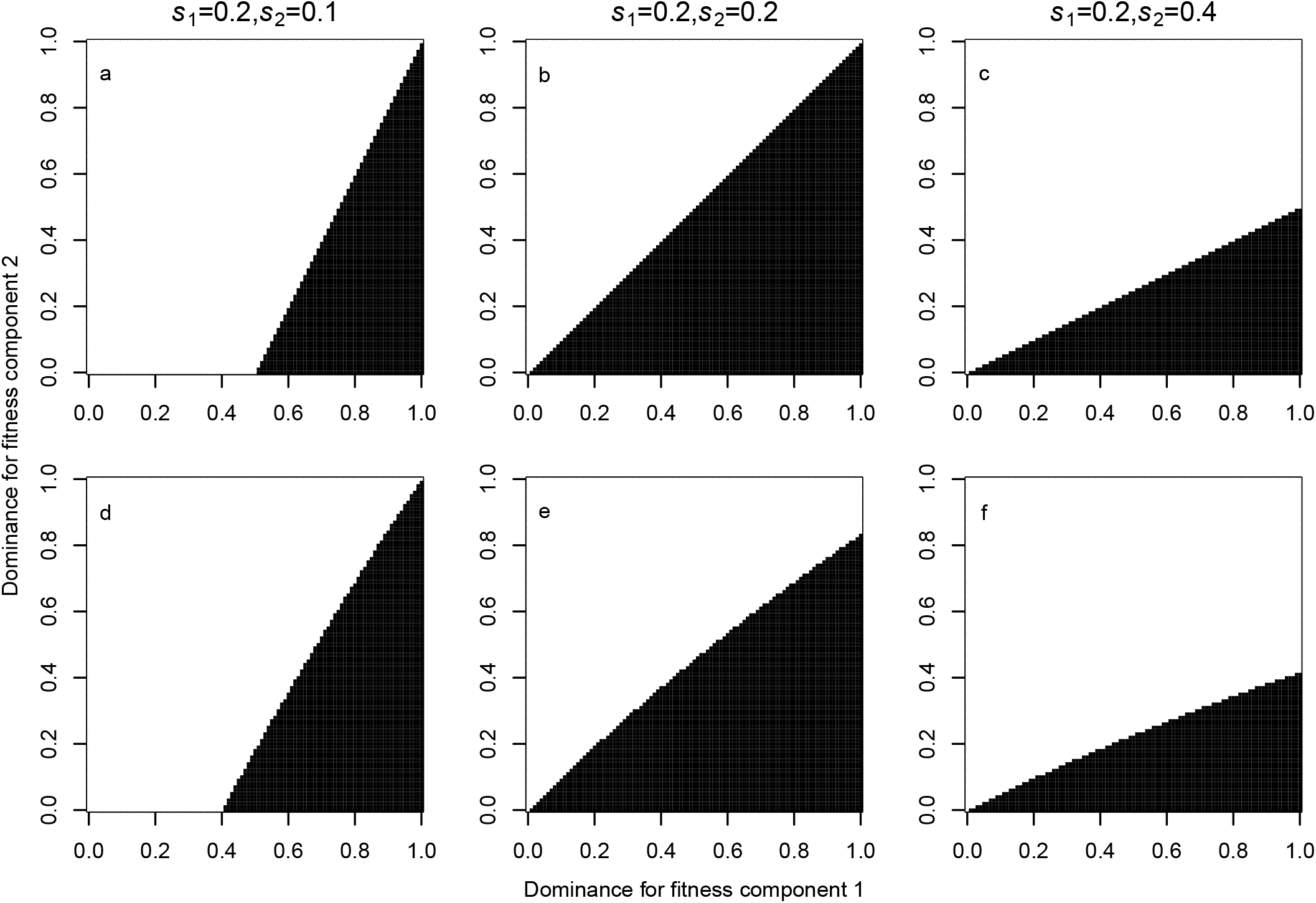
Changes in dominance can maintain polymorphism at a single locus with antagonistic pleiotropy, even without reversal of dominance. The top row shows the region where polymorphism is possible for additive selection and the bottom row for multiplicative selection.

