## Supplemental Code S1 for "How does plant chemodiversity evolve? Testing five hypotheses in one population genetic model": simple_chemodiversity_figures.Rmd

````
---
title: "R code to produce the figures in Wittmann & Bräutigam (2024) How does plant chemodiversity evolve?  Testing five hypotheses in one population genetic model"
author: "Meike Wittmann"
date: "2024-03-26"
output: html_document
---

```{r setup, include=FALSE}
knitr::opts_chunk$set(echo = TRUE)
```

```{r}
library(lattice)
library(viridis)
require(gridExtra)
library(foreach)
```
#Note: The figures can only be generated if you have previously generated the simulation results in the folders for the various simulation experiments.

# Functions
```{r}
## fecundity function modeling the costs of metabolites
f  <- function(z,c){
    return(exp(-c*z))
}

## benefit of metabolites
b  <- function(z,b0,ahalf,s){
return(b0 + (1-b0)/(1+exp(-s*(z-ahalf))))
}

#Geometric mean fitness
geometricmeanfitness  <- function(zb,zf,c,b0,ahalf,s,pi){#2 different zs are needed here because the dominance values can be different for costs and benefits; for genotypes with all homozygotes they are the same though
    return(b(zb,b0,ahalf,s)^pi * f(zf,c))
}

## Invasion analysis
geometricmeangrowthrate  <- function(zb,zf,c,b0,ahalf,s,pi,theta,residentz){#2 different zs are needed here because the dominance values can be different for costs and benefits; for genotypes with all homozygotes they are the same though
    Wmutantwithherbivore <- b(zb,b0,ahalf,s) * f(zf,c)
    Wmutantwithoutherbivore  <- f(zf,c)
    Wresidentwithherbivore  <- b(residentz,b0,ahalf,s) * f(residentz,c)
    Wresidentwithoutherbivore  <- f(residentz,c)
    return((1-theta + theta * Wmutantwithherbivore/Wresidentwithherbivore)^pi * (1-theta + theta * Wmutantwithoutherbivore/Wresidentwithoutherbivore)^(1-pi))
}

## Invasion analysis for generalist-specialist case
geometricmeangrowthrate2herbivores  <- function(zb,zf,c,b0,ahalf,s,p1,p2,theta,residentz,effectsize1,effectsize2){#2 different zs are needed here because the dominance values can be different for costs and benefits; for genotypes with all homozygotes they are the same though
    zb1  <- zb * effectsize1
    zb2  <- zb * effectsize2
    Wmutantwith1and2 <- b(zb1,b0,ahalf,s) * b(zb2,b0,ahalf,s) * f(zf,c)
    Wmutantwith1 <- b(zb1,b0,ahalf,s) * f(zf,c)
    Wmutantwith2 <- b(zb2,b0,ahalf,s) * f(zf,c)
    Wmutantwithoutherbivores  <- f(zf,c)

    m1  <- residentz*effectsize1
    m2  <- residentz*effectsize2
    Wresidentwith1and2  <- b(m1,b0,ahalf,s) *b(m2,b0,ahalf,s) * f(residentz,c)
    Wresidentwith1  <- b(m1,b0,ahalf,s) * f(residentz,c)
    Wresidentwith2  <- b(m2,b0,ahalf,s) * f(residentz,c)
    Wresidentwithoutherbivores  <- f(residentz,c)
    return((1-theta + theta * Wmutantwith1and2/Wresidentwith1and2)^(p1*p2) * (1-theta + theta * Wmutantwith1/Wresidentwith1)^(p1*(1-p2)) * (1-theta + theta * Wmutantwith2/Wresidentwith2)^(p2*(1-p1)) * (1-theta + theta * Wmutantwithoutherbivores/Wresidentwithoutherbivores)^((1-p1)*(1-p2)))
}

geometricmeanfitness2herbivores <- function(zb,zf,c,b0,ahalf,s,p1,p2,effectsize1,effectsize2){
  zb1  <- zb * effectsize1
  zb2  <- zb * effectsize2
  return(b(zb1,b0,ahalf,s)^p1 * b(zb2,b0,ahalf,s)^p2 * f(zf,c))
}

```

# Default values of the parameters
```{r}
# population size
Ndefault <- 500
N <- Ndefault

# adult death probability
thetadefault <- 1 #non-overlapping generations
theta <- thetadefault

# number of generations per phase
gdefault <- 5
g <- gdefault

# probability that a herbivore is present and active during a particular phase
pdefault <- 0.2
p <- pdefault

# number of loci
Ldefault <- 10
L <- Ldefault

# protective effect of a metabolite against a herbivore
mdefault <- 1
m <- mdefault

# baseline probability to escape a herbivore
b0default <- 0
b0 <- b0default

# half-saturation parameter of the benefit function
ahalfdefault <- 0
ahalf <- ahalfdefault

# sensitivity parameter of the benefit function
sdefault <- 1
s <- sdefault

# cost parameter
cdefault <- 0.02
c <- cdefault

# mutation probability per allele copy per generation
udefault <- 0.0001
u <- udefault

# number of replicates for the simulation experiments
numreps <- 5

# number of generations to simulate
numgendefault  <- 5000 #total number of generations to simulate
numgen <- numgendefault
```

# Overview of simulation experiments

Unless otherwise noted, the simulation experiments use the default parameters, iterate through a grid of dominance parameters, and look at the cases of 1, 2 and 5 herbivores.

* chemsim 2: each metabolite active against only 1 herbivore
* chemsim 3: distribution of only repellent effects
* chemsim 4: distribution of repellent and attractive effects
* chemsim 5: p = 1, each metabolite active against only 1 herbivore
* chemsim 6: p = 1, distribution of only repellent effects
* chemsim 7: p = 1, distribution of repellent and attractive effects
* chemsim 8: p = 1, b0 = 0.8, each metabolite active against only 1 herbivore
* chemsim 9: p = 1, b0 = 0.8, distribution of only repellent effects
* chemsim 10: p = 1, b0 = 0.8, distribution of repellent and attractive effects
* chemsim 11: theta = 0.3, each metabolite active against only 1 herbivore
* chemsim 12: theta = 0.3, distribution of only repellent effects
* chemsim 13: theta = 0.3, distribution of repellent and attractive effects
* chemsim 14: p = 1, b0 = 0.8, theta = 0.3, each metabolite active against only 1 herbivore
* chemsim 15: p = 1, b0 = 0.8, theta = 0.3, distribution of only repellent effects
* chemsim 16: p = 1, b0 = 0.8, theta = 0.3, distribution of repellent and attractive effects
* chemsim 17: generalist-specialist scenario with 1 generalist deterred by all metabolites and 1 specialist attracted to all metabolites, uncorrelated occurrence of generalist and specialist
* chemsim 18: generalist-specialist scenario where generalist comes with probability 0.5 and specialist with probability 0.2
* chemsim 19: p = 0.2, b0 = 0, effect size on generalist 0.2, 0.5 or 1, effect size on specialist 0.2
* chemsim 20: p = 1, b0 = 0.8, effect size on generalist 0.2, 0.5 or 1, effect size on specialist 0.2
* chemsim 21: p = 1, b0 = 0, effect size on generalist 0.2, 0.5 or 1, effect size on specialist 0.2
* chemsim 22: rerun of chemsim3, sampling every time step 
* chemsim 23: rerun of chemsim4, sampling every time step
* chemsim 24: zhalf = 5
* chemsim 25: theta = 0.3, zhalf = 5
* chemsim 26: p = 1, zhalf = 5
* chemsim 27: p = 1, theta = 0.3, b0 = 0.8, zhalf = 5
* chemsim 28: p=1, theta = 0.3
* chemsim 29: 1 herbivore, b(0) = 0.01
* chemsim 30: 1 herbivore, p=1, b(0) = 0.01
* chemsim 31: 1 herbivore, p=1, b(0) = 0.8


# Fig 1. Benefit and cost functions
```{r}
par(mfrow=c(1,2))
zvec  <- seq(-5,L,0.01)
plot(zvec,b(zvec,b0,ahalf,s),type="l",xlab=expression(paste("Activity ",italic(a[i]))),ylab=expression(paste("Protection from herbivore ",italic(i))),main=expression(paste("a:  ",italic(a)[half],"=0",sep="")),ylim=c(-0.1,1),col="blue")
b0 <- 0.8
lines(zvec,b(zvec,b0,ahalf,s),col="green")
b0 <- b0default
legend("bottomright",col=c("blue","green"),lty=1,legend=c(0,0.8),title=expression(b[0]))
abline(v=0)
ahalfvec <- c(0,2,4,6)
bofzero <- 0.01
b0vec <- (bofzero * (1+exp(s*ahalfvec)) -1)/exp(s*ahalfvec)
zvec  <- seq(-5,L,0.01)
plot(zvec,b(zvec,b0vec[1],ahalfvec[1],s),type="l",xlab=expression(paste("Activity ",italic(a[i]))),ylab="",main=expression(paste("b:  ",italic(a)[half],">= 0",sep="")),ylim=c(-0.1,1),col=rainbow(4)[1])
lines(zvec,b(zvec,b0vec[2],ahalfvec[2],s),col=rainbow(4)[2])
lines(zvec,b(zvec,b0vec[3],ahalfvec[3],s),col=rainbow(4)[3])
lines(zvec,b(zvec,b0vec[4],ahalfvec[4],s),col=rainbow(4)[4])
abline(v=0)
legend("topleft",legend=ahalfvec,title=expression(italic(a)[half]),col=rainbow(4),lty=1)
text(7,-0.07,"b(0)=0.01")
lines(c(4,4,11),c(-1,0.01,0.01))
dev.print(dev=pdf,"costsandbenefits.pdf")
```

# Plots below Fig. 5
```{r}
pdf("synergyinset.pdf",width=7,height=1.75)
par(mfrow=c(1,4),mar=c(2,2,1,1),oma=c(2,2,0,0))
zvec  <- seq(0,L,0.01)
ahalfvec <- c(0,2,4,6)
bofzero <- 0.01
b0vec <- (bofzero * (1+exp(s*ahalfvec)) -1)/exp(s*ahalfvec)
for (i in 1:4){
  plot(zvec,b(zvec,b0vec[i],ahalfvec[i],s),type="l",ylab="",xlab="",axes=F,ylim=c(0,1),yaxs="i",xaxs="i")
  axis(1)
  axis(2,at=c(0,0.5,1))
}

  title(xlab="Activity",ylab="Protection from herbivore",outer=TRUE,line=0.5)
dev.off()
```


# Fig. 2 Role of dominance and fluctuations with non-overlapping generations
```{r}
pdf("1herbivorenonoverlapping.pdf",width=12/1.5,height=19/1.5)
par(mfcol=c(5,3),mar=c(2,2,3.5,2),oma=c(2,2,0.5,4))
pivec  <- c(0.2,1,1)
b0vec <-c(0,0,0.8)

#ids of the simulation experiments that are used here (see list above)
simid <- c(2,5,8)

#treatments of the simulation experiments
nhtreatments  <- c(1,2,5)
dbtreatments  <- seq(0,1,0.1)
dctreatments  <- seq(0,1,0.1)
NH  <- rep(nhtreatments,length(dbtreatments)*length(dctreatments))
DB  <- rep(rep(dbtreatments,rep(length(nhtreatments),length(dbtreatments))),length(dctreatments))
DC  <- rep(dctreatments,rep(length(dbtreatments)*length(nhtreatments),length(dctreatments)))

for (pit in 1:length(pivec)){
    pi  <- pivec[pit]
    b0  <- b0vec[pit]
    
    zvec  <- seq(0,L,0.01)
    
    plot(zvec,geometricmeanfitness(zvec,zvec,c,b0,ahalf,s,pi),type="l",xlab="",ylab="")

    homozygotez  <- 0:L
    dvec  <- seq(0,1,0.01) #vector of dominance values to be tested
    polymorphismpossible  <- matrix(0,nrow=length(dvec),ncol=length(dvec))

    for (i in 1:length(dvec)){
        for (j in 1:length(dvec)){
            db  <- dvec[i]
            dc  <- dvec[j]
            invasibility  <- logical(length(homozygotez))
            for (k in 1:length(homozygotez)){
                if (homozygotez[k]<L) {
                    largercaninvade  <- geometricmeangrowthrate(homozygotez[k]+db,homozygotez[k]+dc,c,b0,ahalf,s,pi,theta,homozygotez[k])>1
                } else {
                    largercaninvade  <- FALSE
                }
                if (homozygotez[k]>0) {
                    smallercaninvade  <- geometricmeangrowthrate(homozygotez[k]-1+db,homozygotez[k]-1+dc,c,b0,ahalf,s,pi,theta,homozygotez[k])>1
                } else {
                    smallercaninvade  <- FALSE
                }
                if (largercaninvade || smallercaninvade) {
                    invasibility[k]  <- TRUE
                }
            }
            if (sum(invasibility) == length(invasibility)) polymorphismpossible[i,j]  <- 1 #polymorphism is possible if all homozygotes can be invaded
        }
    }

    image(polymorphismpossible,col=c("white","black"),xlab="", ylab="")
    text(0.8,0.05,paste(round(sum(polymorphismpossible)/length(polymorphismpossible)*100,2),"%"),col="white",cex=1.5)
    
    # summarize the simulation results
    setwd(paste("../Simulation/chemsim",simid[pit],sep=""))
    MPI  <- matrix(0,nrow=length(NH)/3,ncol=numreps)
    TMP  <- MPI
    DM  <- MPI
    
    oneherbivoretreatments <- seq(1,length(NH),3)
    for (i in 1:length(oneherbivoretreatments)){ #here we just need the simulations with one herbivore
      for (r in 1:numreps){
        D  <- read.table(paste("output_",oneherbivoretreatments[i],"_",r,"_mutationcounts.txt",sep=""))
        
        ##Allele frequencies
        P  <- D[,-(1:2)]/(2*N)
        
        ##Average number of metabolites per individual
        mpi  <- rowSums(1-(1-P)^2)
        MPI[i,r]  <- mpi[length(mpi)]
        
        ##Total number of metabolites in the population
        tmp  <- rowSums(P>0)
        TMP[i,r]  <- tmp[length(tmp)]
        
        ##Average number of differences
        dm  <- rowSums(2*(1-P)^2 * (1-(1-P)^2))
        DM[i,r]  <- dm[length(dm)]
      }
    }
    
    #trellis.par.set(regions=list(col=plasma(50)))
    
    TMP1  <- matrix(rowMeans(TMP),nrow=11,ncol=11)
    MPI1  <- matrix(rowMeans(MPI),nrow=11,ncol=11)
    DM1  <- matrix(rowMeans(DM),nrow=11,ncol=11)
    print(paste(mean(TMP1),mean(MPI1),mean(DM1)))
    
    image(dbtreatments, dctreatments,TMP1,xlab="",ylab="",main="",breaks=seq(0,10,0.5),col=terrain.colors(20))
    image(dbtreatments, dctreatments,MPI1,xlab="",ylab="",breaks=seq(0,10,0.5),col=terrain.colors(20))
    
    image(dbtreatments, dctreatments,DM1,xlab="",ylab="",breaks=seq(0,10,0.5),col=terrain.colors(20))
    
}

dev.off()
b0 <- b0default
```
Generate color legend
```{r}
pdf("terraincolorbar.pdf",2,8)
image(c(1),seq(0,10,0.5),matrix(seq(0,10,0.5),nrow=1),col=terrain.colors(20),ylab="",xlab="",axes=FALSE)
axis(4,las=1)
dev.off()
```


# Fig. S1: Role of dominance and fluctuations with overlapping generations; values also used in Table S1
```{r}
pdf("1herbivoreoverlapping.pdf",width=12/1.5,height=19/1.5)
par(mfcol=c(5,3),mar=c(2,2,3.5,2),oma=c(2,2,0.5,4))
pivec  <- c(0.2,1,1)
b0vec <-c(0,0,0.8)
theta <- 0.3

#ids of the simulation experiments that are used here (see list above)
simid <- c(11,28,14)

#treatments of the simulation experiments
nhtreatments  <- c(1,2,5)
dbtreatments  <- seq(0,1,0.1)
dctreatments  <- seq(0,1,0.1)
NH  <- rep(nhtreatments,length(dbtreatments)*length(dctreatments))
DB  <- rep(rep(dbtreatments,rep(length(nhtreatments),length(dbtreatments))),length(dctreatments))
DC  <- rep(dctreatments,rep(length(dbtreatments)*length(nhtreatments),length(dctreatments)))

for (pit in 1:length(pivec)){
    pi  <- pivec[pit]
    b0  <- b0vec[pit]
    
    zvec  <- seq(0,L,0.01)
    
    plot(zvec,geometricmeanfitness(zvec,zvec,c,b0,ahalf,s,pi),type="l",xlab="",ylab="")

    homozygotez  <- 0:L
    dvec  <- seq(0,1,0.01) #vector of dominance values to be tested
    polymorphismpossible  <- matrix(0,nrow=length(dvec),ncol=length(dvec))

    for (i in 1:length(dvec)){
        for (j in 1:length(dvec)){
            db  <- dvec[i]
            dc  <- dvec[j]
            invasibility  <- logical(length(homozygotez))
            for (k in 1:length(homozygotez)){
                if (homozygotez[k]<L) {
                    largercaninvade  <- geometricmeangrowthrate(homozygotez[k]+db,homozygotez[k]+dc,c,b0,ahalf,s,pi,theta,homozygotez[k])>1
                } else {
                    largercaninvade  <- FALSE
                }
                if (homozygotez[k]>0) {
                    smallercaninvade  <- geometricmeangrowthrate(homozygotez[k]-1+db,homozygotez[k]-1+dc,c,b0,ahalf,s,pi,theta,homozygotez[k])>1
                } else {
                    smallercaninvade  <- FALSE
                }
                if (largercaninvade || smallercaninvade) {
                    invasibility[k]  <- TRUE
                }
            }
            if (sum(invasibility) == length(invasibility)) polymorphismpossible[i,j]  <- 1 #polymorphism is possible if all homozygotes can be invaded
        }
    }

    image(polymorphismpossible,col=c("white","black"),xlab="", ylab="")
    text(0.8,0.05,paste(round(sum(polymorphismpossible)/length(polymorphismpossible)*100,2),"%"),col="white",cex=1.5)
    
    # summarize the simulation results
    setwd(paste("../Simulation/chemsim",simid[pit],sep=""))
    MPI  <- matrix(0,nrow=length(NH)/3,ncol=numreps)
    TMP  <- MPI
    DM  <- MPI
    
    oneherbivoretreatments <- seq(1,length(NH),3)
    for (i in 1:length(oneherbivoretreatments)){ #here we just need the simulations with one herbivore
      for (r in 1:numreps){
        D  <- read.table(paste("output_",oneherbivoretreatments[i],"_",r,"_mutationcounts.txt",sep=""))
        
        ##Allele frequencies
        P  <- D[,-(1:2)]/(2*N)
        
        ##Average number of metabolites per individual
        mpi  <- rowSums(1-(1-P)^2)
        MPI[i,r]  <- mpi[length(mpi)]
        
        ##Total number of metabolites in the population
        tmp  <- rowSums(P>0)
        TMP[i,r]  <- tmp[length(tmp)]
        
        ##Average number of differences
        dm  <- rowSums(2*(1-P)^2 * (1-(1-P)^2))
        DM[i,r]  <- dm[length(dm)]
      }
    }
    
    #trellis.par.set(regions=list(col=plasma(50)))
    
    TMP1  <- matrix(rowMeans(TMP),nrow=11,ncol=11)
    MPI1  <- matrix(rowMeans(MPI),nrow=11,ncol=11)
    DM1  <- matrix(rowMeans(DM),nrow=11,ncol=11)
    print(paste(mean(TMP1),mean(MPI1),mean(DM1)))
    
    image(dbtreatments, dctreatments,TMP1,xlab="",ylab="",main="",breaks=seq(0,10,0.5),col=terrain.colors(20))
    image(dbtreatments, dctreatments,MPI1,xlab="",ylab="",breaks=seq(0,10,0.5),col=terrain.colors(20))
    
    image(dbtreatments, dctreatments,DM1,xlab="",ylab="",breaks=seq(0,10,0.5),col=terrain.colors(20))
    
}

dev.off()
b0 <- b0default
theta <- thetadefault
```
# Overlapping generations with a smaller death probability per generation (theta = 0.07); Figure not shown, but values used in Table S1
```{r}
pdf("1herbivoreoverlapping_longlife.pdf",width=12/1.5,height=19/1.5)
par(mfcol=c(5,3),mar=c(2,2,3.5,2),oma=c(2,2,0.5,4))
pivec  <- c(0.2,1,1)
b0vec <-c(0,0,0.8)
theta <- 0.07

#ids of the simulation experiments that are used here (see list above)
simid <- c(32,34,33)

#treatments of the simulation experiments
nhtreatments  <- c(1,2,5)
dbtreatments  <- seq(0,1,0.1)
dctreatments  <- seq(0,1,0.1)
NH  <- rep(nhtreatments,length(dbtreatments)*length(dctreatments))
DB  <- rep(rep(dbtreatments,rep(length(nhtreatments),length(dbtreatments))),length(dctreatments))
DC  <- rep(dctreatments,rep(length(dbtreatments)*length(nhtreatments),length(dctreatments)))

for (pit in 1:length(pivec)){
    pi  <- pivec[pit]
    b0  <- b0vec[pit]
    
    zvec  <- seq(0,L,0.01)
    
    plot(zvec,geometricmeanfitness(zvec,zvec,c,b0,ahalf,s,pi),type="l",xlab="",ylab="")

    homozygotez  <- 0:L
    dvec  <- seq(0,1,0.01) #vector of dominance values to be tested
    polymorphismpossible  <- matrix(0,nrow=length(dvec),ncol=length(dvec))

    for (i in 1:length(dvec)){
        for (j in 1:length(dvec)){
            db  <- dvec[i]
            dc  <- dvec[j]
            invasibility  <- logical(length(homozygotez))
            for (k in 1:length(homozygotez)){
                if (homozygotez[k]<L) {
                    largercaninvade  <- geometricmeangrowthrate(homozygotez[k]+db,homozygotez[k]+dc,c,b0,ahalf,s,pi,theta,homozygotez[k])>1
                } else {
                    largercaninvade  <- FALSE
                }
                if (homozygotez[k]>0) {
                    smallercaninvade  <- geometricmeangrowthrate(homozygotez[k]-1+db,homozygotez[k]-1+dc,c,b0,ahalf,s,pi,theta,homozygotez[k])>1
                } else {
                    smallercaninvade  <- FALSE
                }
                if (largercaninvade || smallercaninvade) {
                    invasibility[k]  <- TRUE
                }
            }
            if (sum(invasibility) == length(invasibility)) polymorphismpossible[i,j]  <- 1 #polymorphism is possible if all homozygotes can be invaded
        }
    }

    image(polymorphismpossible,col=c("white","black"),xlab="", ylab="")
    text(0.8,0.05,paste(round(sum(polymorphismpossible)/length(polymorphismpossible)*100,2),"%"),col="white",cex=1.5)
    
    # summarize the simulation results
    setwd(paste("../Simulation/chemsim",simid[pit],sep=""))
    MPI  <- matrix(0,nrow=length(NH)/3,ncol=numreps)
    TMP  <- MPI
    DM  <- MPI
    
    oneherbivoretreatments <- seq(1,length(NH),3)
    for (i in 1:length(oneherbivoretreatments)){ #here we just need the simulations with one herbivore
      for (r in 1:numreps){
        D  <- read.table(paste("output_",oneherbivoretreatments[i],"_",r,"_mutationcounts.txt",sep=""))
        
        ##Allele frequencies
        P  <- D[,-(1:2)]/(2*N)
        
        ##Average number of metabolites per individual
        mpi  <- rowSums(1-(1-P)^2)
        MPI[i,r]  <- mpi[length(mpi)]
        
        ##Total number of metabolites in the population
        tmp  <- rowSums(P>0)
        TMP[i,r]  <- tmp[length(tmp)]
        
        ##Average number of differences
        dm  <- rowSums(2*(1-P)^2 * (1-(1-P)^2))
        DM[i,r]  <- dm[length(dm)]
      }
    }
    
    #trellis.par.set(regions=list(col=plasma(50)))
    
    TMP1  <- matrix(rowMeans(TMP),nrow=11,ncol=11)
    MPI1  <- matrix(rowMeans(MPI),nrow=11,ncol=11)
    DM1  <- matrix(rowMeans(DM),nrow=11,ncol=11)
    print(paste(mean(TMP1),mean(MPI1),mean(DM1)))
    
    image(dbtreatments, dctreatments,TMP1,xlab="",ylab="",main="",breaks=seq(0,10,0.5),col=terrain.colors(20))
    image(dbtreatments, dctreatments,MPI1,xlab="",ylab="",breaks=seq(0,10,0.5),col=terrain.colors(20))
    
    image(dbtreatments, dctreatments,DM1,xlab="",ylab="",breaks=seq(0,10,0.5),col=terrain.colors(20))
    print(mean(DM1))
}

dev.off()
b0 <- b0default
theta <- thetadefault
```

# Fig. 3: Generalist-specialist figure
```{r}
numreps <- 5 #number of replicates
nl <- 10 #number of loci
N <- 500 #population size
g <- 5 #number of generations per phase
numgen  <- 5000 #total number of generations to simulate

## Treatments in all simulation experiments
effectsizespecialist <- - 0.2

#treatments of the simulation experiments
estreatments  <- c(0.2,0.5,1)
dbtreatments  <- seq(0,1,0.1)
dctreatments  <- seq(0,1,0.1)
ES  <- rep(estreatments,length(dbtreatments)*length(dctreatments))
DB  <- rep(rep(dbtreatments,rep(length(estreatments),length(dbtreatments))),length(dctreatments))
DC  <- rep(dctreatments,rep(length(dbtreatments)*length(estreatments),length(dctreatments)))

```

```{r}
simexps <- c(19,21,20)
pivec  <- c(0.2,1,1)
b0vec <-c(0,0,0.8)

MPI_mean <- matrix(0,nrow=3,ncol=length(simexps)) # The 3 is because of 3 different generalist effect sizes
MPI_se <- MPI_mean
TMP_mean <- MPI_mean
TMP_se <-  MPI_mean
DM_mean <- MPI_mean
DM_se <- MPI_mean  
SP <- MPI_mean #scope for polymorphism determined analytically

for (se in 1:length(simexps)){
  
  pi <- pivec[se]
  b0 <- b0vec[se]
  for (est in 1:length(estreatments)){
    effectsizegeneralist <- estreatments[est]
    
    homozygotez  <- 0:L
    dvec  <- seq(0,1,0.01) #vector of dominance values to be tested
    polymorphismpossible  <- matrix(0,nrow=length(dvec),ncol=length(dvec))
    
    for (i in 1:length(dvec)){
      for (j in 1:length(dvec)){
        db  <- dvec[i]
        dc  <- dvec[j]
        invasibility  <- logical(length(homozygotez))
        for (k in 1:length(homozygotez)){
          if (homozygotez[k]<L) {
            largercaninvade  <- geometricmeangrowthrate2herbivores(homozygotez[k]+db,homozygotez[k]+dc,c,b0,ahalf,s,pi,pi,theta,homozygotez[k],effectsizegeneralist,effectsizespecialist)>1
          } else {
            largercaninvade  <- FALSE
          }
          if (homozygotez[k]>0) {
            smallercaninvade  <- geometricmeangrowthrate2herbivores(homozygotez[k]-1+db,homozygotez[k]-1+dc,c,b0,ahalf,s,pi,pi,theta,homozygotez[k],effectsizegeneralist,effectsizespecialist)>1
          } else {
            smallercaninvade  <- FALSE
          }
          if (largercaninvade || smallercaninvade) {
            invasibility[k]  <- TRUE
          }
        }
        if (sum(invasibility) == length(invasibility)) polymorphismpossible[i,j]  <- 1 #polymorphism is possible if all homozygotes can be invaded
      }
    }
    
    SP[est,se] <- sum(polymorphismpossible)/length(polymorphismpossible)*100
    
  }
  
  setwd(paste("../Simulation/chemsim",simexps[se],sep=""))
  MPI  <- matrix(0,nrow=length(ES),ncol=numreps)
  TMP  <- MPI
  DM  <- MPI
  
  for (i in 1:length(ES)){
    for (r in 1:numreps){
      D  <- read.table(paste("output_",i,"_",r,"_mutationcounts.txt",sep=""))
      
      ##Allele frequencies
      P  <- D[,-(1:2)]/(2*N)
      
      ##Average number of metabolites per individual
      mpi  <- rowSums(1-(1-P)^2)
      MPI[i,r]  <- mpi[length(mpi)]
      
      ##Total number of metabolites in the population
      tmp  <- rowSums(P>0)
      TMP[i,r]  <- tmp[length(tmp)]
      
      ##Average number of differences
      dm  <- rowSums(2*(1-P)^2 * (1-(1-P)^2))
      DM[i,r]  <- dm[length(dm)]
    }
  }
  for (nhi in 1:3){
    subMPI <- MPI[seq(nhi,length(ES),3),]
    MPI_mean[nhi,se] <- mean(subMPI)
    MPI_se[nhi,se] <- sd(colMeans(subMPI))/sqrt(numreps)
    
    subTMP <- TMP[seq(nhi,length(ES),3),]
    TMP_mean[nhi,se] <- mean(subTMP)
    TMP_se[nhi,se] <- sd(colMeans(subTMP))/sqrt(numreps)
    
    subDM <- DM[seq(nhi,length(ES),3),]
    DM_mean[nhi,se] <- mean(subDM)
    DM_se[nhi,se] <- sd(colMeans(subDM))/sqrt(numreps)
  }
}

b0 <- b0default
```

```{r}
pdf("summaryfigure_generalist_specialist.pdf",width=8.2,height=3.2)
par(mfrow=c(1,4))
par(mar=c(1,3,2,0.5),oma=c(2.5,1,0.5,1),mgp=c(3,0.6,0))
#encode fluctuation scenario as color
colorvec <- c("blue","magenta","green")
#blue: fluctuations with p=0.2 and b0 = 0, magenta: no fluctuations with p=1 and b0=0, green: no fluctuations with p=1 and b0=0.8

matplot(estreatments,TMP_mean,axes=F,xlab="",ylab="",col=colorvec,pch=15,ylim=c(4,10),main="a",lty=1,type="b")
for (i in 1:length(estreatments)){
  for (j in 1:length(simexps)){
    lines(rep(estreatments[i],2),c(TMP_mean[i,j]-TMP_se[i,j],TMP_mean[i,j]+TMP_se[i,j]),col=colorvec[j])
  }
}
axis(1,at=estreatments,labels=estreatments)
axis(2)
box()
title(ylab="Total number of metabolites in the population",line=1.7)
legend("topleft",col=c("blue","magenta","green"),legend=c("fluctuating herbivory","constant high herbivory","constant low herbivory"),lty=1,lwd=1,pch=15,bty="n")

matplot(estreatments,MPI_mean,axes=F,xlab="",ylab="",col=colorvec,pch=15,ylim=c(0,10),main="b",lty=1,type="b")
for (i in 1:length(estreatments)){
  for (j in 1:length(simexps)){
    lines(rep(estreatments[i],2),c(MPI_mean[i,j]-MPI_se[i,j],MPI_mean[i,j]+MPI_se[i,j]),col=colorvec[j])
  }
}
axis(1,at=estreatments,labels=estreatments)
axis(2)
box()
title(ylab="Average number of metabolites per individual",line=1.7)

matplot(estreatments,DM_mean,axes=F,xlab="",ylab="",col=colorvec,pch=15,ylim=c(0,2),main="c",lty=1,type="b")
for (i in 1:length(estreatments)){
  for (j in 1:length(simexps)){
    lines(rep(estreatments[i],2),c(DM_mean[i,j]-DM_se[i,j],DM_mean[i,j]+DM_se[i,j]),col=colorvec[j])
  }
}
axis(1,at=estreatments,labels=estreatments)
axis(2)
box()
title(ylab="Average number of differences between individuals",line=1.7)

matplot(estreatments,SP,axes=F,xlab="",ylab="",col=colorvec,pch=15,ylim=c(0,100),main="d",lty=1,type="b")
axis(1,at=estreatments,labels=estreatments)
axis(2)
box()
title(ylab="Scope for polymorphism in %  (analytic)",line=1.7)

title(xlab="Protective effect of metabolites against generalist",outer=TRUE,line=1)
dev.off()
```
 
# Fig. 4: Role of number of herbivores

Set common parameters
```{r}
numreps <- 5 #number of replicates
nl <- 10 #number of loci
N <- 500 #population size
g <- 5 #number of generations per phase
numgen  <- 5000 #total number of generations to simulate

## Treatments in all simulation experiments
nhtreatments  <- c(1,2,5)
dbtreatments  <- seq(0,1,0.1)
dctreatments  <- seq(0,1,0.1)
NH  <- rep(nhtreatments,length(dbtreatments)*length(dctreatments))
DB  <- rep(rep(dbtreatments,rep(length(nhtreatments),length(dbtreatments))),length(dctreatments))
DC  <- rep(dctreatments,rep(length(dbtreatments)*length(nhtreatments),length(dctreatments)))
NH
```

Use time point 49 as a reference for all because time point 50 might already be affected by invasions in some of the experiments.
```{r}
referencetime <- 49
```

```{r}
simexps <- c(3,4,6,7,9,10)
MPI_mean <- matrix(0,nrow=3,ncol=length(simexps)) # The 3 is because of three different herbivore number treatments
MPI_se <- MPI_mean
TMP_mean <- MPI_mean
TMP_se <-  MPI_mean
DM_mean <- MPI_mean
DM_se <- MPI_mean  
  
for (se in 1:length(simexps)){
  setwd(paste("../Simulation/chemsim",simexps[se],sep=""))
  MPI  <- matrix(0,nrow=length(NH),ncol=numreps)
  TMP  <- MPI
  DM  <- MPI
  for (i in 1:length(NH)){
    for (r in 1:numreps){
      D  <- read.table(paste("output_",i,"_",r,"_mutationcounts.txt",sep=""))
      
      ##Allele frequencies
      P  <- D[,-(1:2)]/(2*N)
      
      ##Average number of metabolites per individual
      mpi  <- rowSums(1-(1-P)^2)
      MPI[i,r]  <- mpi[referencetime]
      
      ##Total number of metabolites in the population
      tmp  <- rowSums(P>0)
      TMP[i,r]  <- tmp[referencetime]
      
      ##Average number of differences
      dm  <- rowSums(2*(1-P)^2 * (1-(1-P)^2))
      DM[i,r]  <- dm[referencetime]
    }
  }
  for (nhi in 1:3){
    subMPI <- MPI[seq(nhi,length(NH),3),]
    MPI_mean[nhi,se] <- mean(subMPI)
    MPI_se[nhi,se] <- sd(colMeans(subMPI))/sqrt(numreps)
    
    subTMP <- TMP[seq(nhi,length(NH),3),]
    TMP_mean[nhi,se] <- mean(subTMP)
    TMP_se[nhi,se] <- sd(colMeans(subTMP))/sqrt(numreps)
    
    subDM <- DM[seq(nhi,length(NH),3),]
    DM_mean[nhi,se] <- mean(subDM)
    DM_se[nhi,se] <- sd(colMeans(subDM))/sqrt(numreps)
  }
}
```
```{r}
TMP_mean
```

```{r}
pdf("summaryfigure.pdf",width=7.5,height=3.2)
par(mfrow=c(1,3))
par(mar=c(1,3,2,0.5),oma=c(2.5,1,0.5,1),mgp=c(3,0.6,0))
TMP_mean_vec <- as.numeric(TMP_mean)
TMP_se_vec <- as.numeric(TMP_se)
locationvec <- rep(c(1,2,5),length(simexps))# +runif(length(TMP_mean_vec),-0.1,0.1)
#encode fluctuation scenario as color
colorvec <- c(rep("blue",2*3),rep("magenta",2*3),rep("green",2*3))
colorvec2 <- c(rep("blue",2),rep("magenta",2),rep("green",2))
#blue: fluctuations with p=0.2 and b0 = 0, magenta: no fluctuations with p=1 and b0=0, green: no fluctuations with p=1 and b0=0.8
pchvec <- rep(c(16,17),3)
matplot(c(1,2,5),TMP_mean,axes=F,xlab="",ylab="",col=colorvec2,pch=pchvec,ylim=c(4,10),main="a",lty=1,type="b")
for (i in 1:length(locationvec)){
  lines(rep(locationvec[i],2),c(TMP_mean_vec[i]-TMP_se_vec[i],TMP_mean_vec[i]+TMP_se_vec[i]),col=colorvec[i])
}
axis(1,at=c(1,2,5),labels=c("1", "2", "5"))
axis(2)
box()
title(ylab="Total number of metabolites in the population",line=1.7)
legend("bottomleft",col=c("blue","magenta","green"),legend=c("fluctuating herbivory","constant high herbivory","constant low herbivory"),lty=1,lwd=3,bty="n")
legend("topleft",pch=c(16:17,0),legend=c("repell","repell + attract"),bty="n")

MPI_mean_vec <- as.numeric(MPI_mean)
MPI_se_vec <- as.numeric(MPI_se)
matplot(c(1,2,5),MPI_mean,axes=F,ylab="",xlab="",col=colorvec2,pch=pchvec,ylim=c(0,10),main="b",lty=1,type="b")
for (i in 1:length(locationvec)){
  lines(rep(locationvec[i],2),c(MPI_mean_vec[i]-MPI_se_vec[i],MPI_mean_vec[i]+MPI_se_vec[i]),col=colorvec[i])
}
axis(1,at=c(1,2,5),labels=c("1", "2", "5"))
axis(2)
box()
title(ylab="Average number of metabolites per individual", line=1.7)

DM_mean_vec <- as.numeric(DM_mean)
DM_se_vec <- as.numeric(DM_se)
matplot(c(1,2,5),DM_mean,axes=F,xlab="",ylab="",col=colorvec2,pch=pchvec,main="c",ylim=c(0,1.2),lty=1,type="b")
for (i in 1:length(locationvec)){
  lines(rep(locationvec[i],2),c(DM_mean_vec[i]-DM_se_vec[i],DM_mean_vec[i]+DM_se_vec[i]),col=colorvec[i])
}
axis(1,at=c(1,2,5),labels=c("1", "2", "5"))
axis(2,at=seq(0,2,0.5))
box()
title(ylab="Average number of differences between individuals",line=1.7)

title(xlab="Number of herbivores",outer=TRUE,line=1)
dev.off()
```


# Fig. 5 Synergy figure

```{r}
numreps <- 5 #number of replicates
nl <- 10 #number of loci
N <- 500 #population size
g <- 5 #number of generations per phase
numgen  <- 5000 #total number of generations to simulate

## Treatments in all simulation experiments
ahalfvec <- c(0,2,4,6)

pivec <-c(0.2,1,1)

dbtreatments  <- seq(0,1,0.1)
dctreatments  <- seq(0,1,0.1)

AH  <- rep(ahalfvec,length(dbtreatments)*length(dctreatments))
DB  <- rep(rep(dbtreatments,rep(length(ahalfvec),length(dbtreatments))),length(dctreatments))
DC  <- rep(dctreatments,rep(length(dbtreatments)*length(ahalfvec),length(dctreatments)))
```

```{r}
simexps <- c(29:31)
bofzerovec <- c(0.01,0.01,0.8)
MPI_mean <- matrix(0,nrow=4,ncol=length(simexps)) # The 4 is because of four different benefit function treatments
MPI_se <- MPI_mean
TMP_mean <- MPI_mean
TMP_se <-  MPI_mean
DM_mean <- MPI_mean
DM_se <- MPI_mean  
SP <- MPI_mean
  
for (se in 1:length(simexps)){
  
  pi <- pivec[se]
  b0vec <- (bofzerovec[se] * (1+exp(s*ahalfvec)) -1)/exp(s*ahalfvec)
 
  for (ahalft in 1:length(ahalfvec)){
    ahalf <- ahalfvec[ahalft]
    b0 <- b0vec[ahalft]
     
    homozygotez  <- 0:L
    dvec  <- seq(0,1,0.01) #vector of dominance values to be tested
    polymorphismpossible  <- matrix(0,nrow=length(dvec),ncol=length(dvec))
    
    for (i in 1:length(dvec)){
      for (j in 1:length(dvec)){
        db  <- dvec[i]
        dc  <- dvec[j]
        invasibility  <- logical(length(homozygotez))
        for (k in 1:length(homozygotez)){
          if (homozygotez[k]<L) {
            largercaninvade  <- geometricmeangrowthrate(homozygotez[k]+db,homozygotez[k]+dc,c,b0,ahalf,s,pi,theta,homozygotez[k])>1
          } else {
            largercaninvade  <- FALSE
          }
          if (homozygotez[k]>0) {
            smallercaninvade  <- geometricmeangrowthrate(homozygotez[k]-1+db,homozygotez[k]-1+dc,c,b0,ahalf,s,pi,theta,homozygotez[k])>1
          } else {
            smallercaninvade  <- FALSE
          }
          if (largercaninvade || smallercaninvade) {
            invasibility[k]  <- TRUE
          }
        }
        if (sum(invasibility) == length(invasibility)) polymorphismpossible[i,j]  <- 1 #polymorphism is possible if all homozygotes can be invaded
      }
    }
    
    SP[ahalft,se] <- sum(polymorphismpossible)/length(polymorphismpossible)*100
  }
}
  
for (se in 1:length(simexps)){
  
  setwd(paste("../Simulation/chemsim",simexps[se],sep=""))
  MPI  <- matrix(0,nrow=length(AH),ncol=numreps)
  TMP  <- MPI
  DM  <- MPI
  for (i in 1:length(AH)){
    for (r in 1:numreps){
      D  <- read.table(paste("output_",i,"_",r,"_mutationcounts.txt",sep=""))
      
      ##Allele frequencies
      P  <- D[,-(1:2)]/(2*N)
      
      ##Average number of metabolites per individual
      mpi  <- rowSums(1-(1-P)^2)
      MPI[i,r]  <- mpi[length(mpi)]
      
      ##Total number of metabolites in the population
      tmp  <- rowSums(P>0)
      TMP[i,r]  <- tmp[length(tmp)]
      
      ##Average number of differences
      dm  <- rowSums(2*(1-P)^2 * (1-(1-P)^2))
      DM[i,r]  <- dm[length(dm)]
    }
  }
  for (nhi in 1:4){
    subMPI <- MPI[seq(nhi,length(AH),4),]
    MPI_mean[nhi,se] <- mean(subMPI)
    MPI_se[nhi,se] <- sd(colMeans(subMPI))/sqrt(numreps)
    
    subTMP <- TMP[seq(nhi,length(AH),4),]
    TMP_mean[nhi,se] <- mean(subTMP)
    TMP_se[nhi,se] <- sd(colMeans(subTMP))/sqrt(numreps)
    
    subDM <- DM[seq(nhi,length(AH),4),]
    DM_mean[nhi,se] <- mean(subDM)

    DM_se[nhi,se] <- sd(colMeans(subDM))/sqrt(numreps)
  }
  }
  ahalf <- ahalfdefault
```


```{r}
pdf("summaryfigure_synergy.pdf",width=8.2,height=3.2)
par(mfrow=c(1,4))
par(mar=c(1,3,2,0.5),oma=c(2.5,1,0.5,1),mgp=c(3,0.6,0))
#encode fluctuation scenario as color
colorvec <- c("blue","magenta","green")
#blue: fluctuations with p=0.2 and b0 = 0, magenta: no fluctuations with p=1 and b0=0, green: no fluctuations with p=1 and b0=0.8

matplot(ahalfvec,TMP_mean,axes=F,xlab="",ylab="",col=colorvec,pch=15,ylim=c(4,10),main="a",lty=1,type="b")
for (i in 1:length(ahalfvec)){
  for (j in 1:length(simexps)){
    lines(rep(ahalfvec[i],2),c(TMP_mean[i,j]-TMP_se[i,j],TMP_mean[i,j]+TMP_se[i,j]),col=colorvec[j])
  }
}
axis(1,at=seq(0,6,2),labels=c("0", "2", "4", "6"))
axis(2)
box()
legend("bottomleft",col=c("blue","magenta","green"),legend=c("p=0.2, b(0)=0.01","p=1, b(0)=0.01","p=1, b(0)=0.8"),lty=1,lwd=1,pch=15,bty="n")
title(ylab="Total number of metabolites in the population",line=1.7)

matplot(ahalfvec,MPI_mean,axes=F,xlab="",ylab="",col=colorvec,pch=15,ylim=c(0,10),main="b",lty=1,type="b")
for (i in 1:length(ahalfvec)){
  for (j in 1:length(simexps)){
    lines(rep(ahalfvec[i],2),c(MPI_mean[i,j]-MPI_se[i,j],MPI_mean[i,j]+MPI_se[i,j]),col=colorvec[j])
  }
}
axis(1,at=seq(0,6,2),labels=c("0", "2", "4", "6"))
axis(2)
box()
title(ylab="Average number of metabolites per individual",line=1.7)

matplot(ahalfvec,DM_mean,axes=F,xlab="",ylab="",col=colorvec,pch=15,ylim=c(0,2),main="c",lty=1,type="b")
for (i in 1:length(ahalfvec)){
  for (j in 1:length(simexps)){
    lines(rep(ahalfvec[i],2),c(DM_mean[i,j]-DM_se[i,j],DM_mean[i,j]+DM_se[i,j]),col=colorvec[j])
  }
}
axis(1,at=seq(0,6,2),labels=c("0", "2", "4", "6"))
axis(2)
box()
title(ylab="Average number of differences between individuals",line=1.7)

matplot(ahalfvec,SP,axes=F,xlab="",ylab="",col=colorvec,pch=15,ylim=c(0,100),main="d",lty=1,type="b")
axis(1,at=seq(0,6,2),labels=c("0", "2", "4", "6"))
axis(2)
box()
title(ylab="Scope for polymorphism in %  (analytic)",line=1.7)

title(xlab="Half-saturation parameter of the benefit function",outer=TRUE,line=1)
dev.off()
```

Find out why scope of polymorphism is so low for large ahalf in the constant low herbivory scenario.
```{r}
pdf("geometricmeaninset.pdf",width=3,height=2)
par(mar=c(3,3,0,0))
ahalf <- 6
bofzero <- 0.8
pi <- 1
b0 <- (bofzero * (1+exp(s*ahalf)) -1)/exp(s*ahalf)
homozygotez  <- 0:L
homozygotegeometricmean  <- geometricmeanfitness(homozygotez,homozygotez,c,b0,ahalf,s,pi)
print(homozygotegeometricmean)
ahalf <- ahalfdefault
b0 <- b0default
plot(homozygotez,homozygotegeometricmean,type="l",xlab="",ylab="",col="green")
title(xlab="Number of metabolites",ylab="Geom. mean fitness    ",line=2)
```

## Fig. 6 - Fitness after an invasion
Simulation experiments with invasion:
```{r}
invasion_simexps <- c(22,23)
averagefitnessafterinvasion <- matrix(0,nrow=length(invasion_simexps),ncol=length(NH))
MPI <- matrix(0,nrow=length(invasion_simexps),ncol=length(NH))
TMP <- matrix(0,nrow=length(invasion_simexps),ncol=length(NH))
DM <- matrix(0,nrow=length(invasion_simexps),ncol=length(NH))
```

```{r}
for (se in 1:length(invasion_simexps)){
  for (i in 1:length(NH)){
    for (r in 1:numreps){
      fitnesstable  <- read.table(paste("../Simulation/chemsim",invasion_simexps[se],"/output_",i,"_",r,"_fitness.txt",sep=""))
      averagefitness <- fitnesstable[,3]*exp(fitnesstable[,5])
      averagefitnessafterinvasion[se,i] <- averagefitnessafterinvasion[se,i] + mean(averagefitness[-(1:20)])/numreps
      #plot(averagefitness)
      
      D  <- read.table(paste("../Simulation/chemsim",invasion_simexps[se],"/output_",i,"_",r,"_mutationcounts.txt",sep=""))
      
      ##Allele frequencies
      P  <- D[,-(1:2)]/(2*N)
      
      ##Average number of metabolites per individual
      mpi  <- rowSums(1-(1-P)^2)
      MPI[se,i]  <- MPI[se,i] + mean(mpi[1:19])/numreps
      
      ##Total number of metabolites in the population
      tmp  <- rowSums(P>0)
      TMP[se,i]  <- TMP[se,i] + mean(tmp[1:19])/numreps
      
      ##Average number of differences
      dm  <- rowSums(2*(1-P)^2 * (1-(1-P)^2))
      DM[se,i]  <- DM[se,i] + mean(dm[1:19])/numreps
    }
  }
}

```

```{r fig.height=5, fig.width=6}
par(mfrow=c(2,3))
plot(TMP[1,],averagefitnessafterinvasion[1,],xlab="Total metabolites",ylab="Average fitness after invasion",main="(a)",col=1:3,xlim=c(0,10))
legend("topleft",pch=1,col=1:3,legend=c("1 -> 2","2 -> 3","5 -> 6"))
plot(MPI[1,],averagefitnessafterinvasion[1,],xlab="Average number per individual",ylab="Average fitness after invasion",main="(b)",col=1:3,xlim=c(0,10))
plot(DM[1,],averagefitnessafterinvasion[1,],xlab="Average number of differences",ylab="Average fitness after invasion",main="(c)",col=1:3,xlim=c(0,10))
plot(TMP[2,],averagefitnessafterinvasion[2,],xlab="Total metabolites",ylab="Average fitness after invasion",main="(d)",col=1:3,xlim=c(0,10))
plot(MPI[2,],averagefitnessafterinvasion[2,],xlab="Average number per individual",ylab="Average fitness after invasion",main="(e)",col=1:3,xlim=c(0,10))
plot(DM[2,],averagefitnessafterinvasion[2,],xlab="Average number of differences",ylab="Average fitness after invasion",main="(f)",col=1:3,xlim=c(0,10))
dev.print(dev=pdf,"invasionsummary.pdf")
```

# Get loss probability for loci that have at least one attractive effect (reported in the text)
```{r}
simid <- 10
setwd(paste("../Simulation/chemsim",simid,sep=""))
nhtreatments  <- c(1,2,5)
dbtreatments  <- seq(0,1,0.1)
dctreatments  <- seq(0,1,0.1)
NH  <- rep(nhtreatments,length(dbtreatments)*length(dctreatments))
DB  <- rep(rep(dbtreatments,rep(length(nhtreatments),length(dbtreatments))),length(dctreatments))
DC  <- rep(dctreatments,rep(length(dbtreatments)*length(nhtreatments),length(dctreatments)))
referencetime <- 49

par(mfrow=c(1,3))

combinedlossprobs <- foreach(pit=1:length(nhtreatments))%do%{
    oneeffectsizetreatments <- seq(pit,length(NH),3)
    numherbivores <- nhtreatments[pit]
    
    alllossprobs <- foreach (i=1:length(oneeffectsizetreatments),.combine='rbind')%do%{ 
      lossprobsreps <- foreach (r=1:numreps,.combine='c')%do%{
        D  <- read.table(paste("output_",oneeffectsizetreatments[i],"_",r,"_mutationcounts.txt",sep=""))
        M <- read.table(paste("output_",oneeffectsizetreatments[i],"_",r,"_activitymatrix.txt",sep=""))
        
        #Get loci that have at least one attractive effect
        attractiveloci <- which(apply(M,1,min) < 0)
        
        ##Allele frequencies
        P  <- as.numeric(D[referencetime,-(1:2)]/(2*N))
        
        #loss probability of attractive loci
        lossprob <- sum(P[attractiveloci] == 0)/length(attractiveloci)
        return(lossprob)
      }
      return(lossprobsreps)
    }
}
```
```{r}
mean(combinedlossprobs[[1]],na.rm=T)
mean(combinedlossprobs[[2]],na.rm=T)
mean(combinedlossprobs[[3]],na.rm=T)

1-0.9^c(1,2,5)
```

#Fig. S2
Get final allele frequencies for generalist-specialist scenarios and generate a site frequency spectrum
```{r}
effectsizespecialist <- - 0.2
generalisteffectsizevec <- c(0.2,0.5,1)
p = 1
pi = p

#ids of the simulation experiments that are used here (see list above)
simid <- c(19,21,20)
mainvec <- c("a: fluctuating herbivory","b: constant high herbivory", "c: constant low herbivory")
pdf("generalistspecialist_sfs.pdf",height=8,width=6)
par(mfrow=c(3,1))


for (se in 1:3){
setwd(paste("../Simulation/chemsim",simid[se],sep=""))

#treatments of the simulation experiments
nhtreatments  <- c(0.2,5,1)
dbtreatments  <- seq(0,1,0.1)
dctreatments  <- seq(0,1,0.1)
NH  <- rep(nhtreatments,length(dbtreatments)*length(dctreatments))
DB  <- rep(rep(dbtreatments,rep(length(nhtreatments),length(dbtreatments))),length(dctreatments))
DC  <- rep(dctreatments,rep(length(dbtreatments)*length(nhtreatments),length(dctreatments)))

allmeansfs <- matrix(nrow=3,ncol=7)

for (pit in 1:length(generalisteffectsizevec)){
  print(generalisteffectsizevec[pit])
    meanallelefrequencies <- matrix(0,nrow=length(NH)/3,ncol=numreps)
    sdallelefrequencies <- matrix(0,nrow=length(NH)/3,ncol=numreps)
    sfs <- matrix(0,nrow=length(NH)/3,ncol=7)
    
    oneeffectsizetreatments <- seq(pit,length(NH),3)
    for (i in 1:length(oneeffectsizetreatments)){ 
      for (r in 1:numreps){
        D  <- read.table(paste("output_",oneeffectsizetreatments[i],"_",r,"_mutationcounts.txt",sep=""))
        
        ##Allele frequencies
        P  <- D[nrow(D),-(1:2)]/(2*N)
        meanallelefrequencies[i,r] <- mean(as.numeric(P))
        sdallelefrequencies[i,r] <- sd(as.numeric(P))
        
        sfs[i,1] <- sfs[i,1] + sum(P == 0)
        sfs[i,2] <- sfs[i,2] + sum(P > 0 & P < 0.01)
        sfs[i,3] <- sfs[i,3] + sum(P >= 0.01 & P < 0.1)
        sfs[i,4] <- sfs[i,4] + sum(P >= 0.1 & P < 0.9)
        sfs[i,5] <- sfs[i,5] + sum(P >= 0.9 & P < 0.99)
        sfs[i,6] <- sfs[i,6] + sum(P >= 0.99 & P < 1)
        sfs[i,7] <- sfs[i,7] + sum(P == 1)
      }
    }
    sfs <- sfs/numreps
    meansfs <- colMeans(sfs)
    print(meansfs)
    print(mean(meanallelefrequencies))
    print(mean(sdallelefrequencies))
    allmeansfs[pit,] <- meansfs
}

barplot(allmeansfs,beside=T,ylab="Number of loci",col=gray.colors(3),names.arg=c("0","(0,0.01)","[0.01,0.1)","[0.1,0.9)","[0.9,0.99)","[0.99,1)","1"),xlab="Allele frequency",main=mainvec[se])
legend("topright",col=gray.colors(3),title="Protective effect against generalist",legend=c(0.2,0.5,1),pch=15)
}
dev.off()
p <- pdefault
```

#Fig. S4
Comparison to a simple antagonistic pleiotropy model at a single locus as in Rose (1982), but parameterized more like in our model
```{r}
s1 <- 0.2
s2vec <- c(0.1,0.2,0.4)

dvec  <- seq(0,1,0.01) #vector of dominance values to be tested
pdf("Rosemodel.pdf",width=7,height=5)
par(mfcol=c(2,3),mar=c(2,2,2,2),oma=c(2,2,2,0))
labelvec <- c("a","b","c","d","e","f")
mainvec<-c(expression(paste(italic(s[1]),"=0.2,", italic(s[2]),"=0.1")),expression(paste(italic(s[1]),"=0.2,", italic(s[2]),"=0.2")),expression(paste(italic(s[1]),"=0.2,", italic(s[2]),"=0.4")))

for (si in 1:length(s2vec)){
  s2 <- s2vec[si]
  polymorphismpossibleadditive  <- matrix(0,nrow=length(dvec),ncol=length(dvec))
  polymorphismpossiblemultiplicative  <- matrix(0,nrow=length(dvec),ncol=length(dvec))
  
  for (i in 1:length(dvec)){
    for (j in 1:length(dvec)){
      d1  <- dvec[i]
      d2  <- dvec[j]
      polymorphismpossibleadditive[i,j] <- 2+d1*s1-d2*s2 > max(2,2+s1-s2)
      polymorphismpossiblemultiplicative[i,j] <- (1+d1*s1)*(1-d2*s2) > max(1,(1+s1)*(1-s2))
    }
  }
  
  image(polymorphismpossibleadditive,col=c("white","black"),xlab="", ylab="",main=mainvec[si])
  text(0.05,0.9,labelvec[si])
  image(polymorphismpossiblemultiplicative,col=c("white","black"),xlab="", ylab="")
  text(0.05,0.9,labelvec[3+si])
}
title(xlab="Dominance for fitness component 1",ylab="Dominance for fitness component 2",outer=T,line=0.5)
dev.off()
```
````
