## Supplemental Code S1 for "How does plant chemodiversity evolve? Testing five hypotheses in one population genetic model": README.txt

The Simulation folder contains the C++ source code for the simulations in Wittmann & Bräutigam (2024) How does plant chemodiversity evolve?  Testing five hypotheses in one population genetic model, which is derived from simulation code for Wittmann et al. (2017) Segregation lift: How seasonally fluctuating selection can maintain polymorphism at many loci. It is under the GNU General Public License (see COPYING.txt). However, the program should be seen mainly as a reference for the present paper (it is not very user friendly). Note that the simulation program includes some options that are not used in the present paper. To re-compile the simulation program, you need to have the boost C++ libraries installed.

A compiled linux executable is included.

To re-run simulation experiments, place the program executable into the respective simulation experiment folders chemsim2 to chemsim31 and use the script start_simulations.R to start the simulations, ideally on a computer cluster (the scripts will try to parallelize over 20 cores).

The parameter settings of all simulations experiments are explained in the R Markdown file in the RCode folder, which also contains all the code to analyze the simulated data and to generate the figures.
