## Supplemental Code S1 for "How does plant chemodiversity evolve? Testing five hypotheses in one population genetic model": start_simulations.R

```
#!/usr/bin/Rscript --vanilla
args <- commandArgs(TRUE)

library(foreach)
library(doMC)


numreps <- 5
numcores <- 20
registerDoMC(numcores)

#parameters
inputfilenamebase <- "input"
outputfilenamebase <- "output"
allinfofilenamebase <- "allinfo"
fixedfilenamebase <- "fixedfile"
resultfilebase <- "results"

nl <- 10 #number of loci
N <- 500 #population size
g <- 5 #number of generations per phase
numgen  <- 5000 #total number of generations to simulate

pi  <- 0.5
b0  <- 0
zhalf  <- 0
senspar  <- 1
c  <- 0.02
adultdeathrate  <- 1
numherbivores  <- 2 #1 generalist, one specialist


nhtreatments  <- c(0.2,5,1) #variation in effect size
dbtreatments  <- seq(0,1,0.1)
dctreatments  <- seq(0,1,0.1)

NH  <- rep(nhtreatments,length(dbtreatments)*length(dctreatments))
DB  <- rep(rep(dbtreatments,rep(length(nhtreatments),length(dbtreatments))),length(dctreatments))
DC  <- rep(dctreatments,rep(length(dbtreatments)*length(nhtreatments),length(dctreatments)))

foreach (treatment=1:length(NH)) %dopar%{
    effectsize  <- NH[treatment]
    
    foreach (replicate=1:numreps) %do% {
       
        inputfilename <- paste(inputfilenamebase,"_",treatment,"_",replicate,".txt",sep="")
        sink(file=inputfilename)
        cat(paste(outputfilenamebase,"_",treatment,"_",replicate," #filenamebase\n",sep=""))
        cat(paste((replicate-1)*length(nhtreatments)+treatment,"#seed\n"))
        cat(paste(1,"#numreps\n"))
        cat(paste(g,"#generations per phase\n"))
        cat(paste(numgen,"#total number of generations\n"))
        cat(paste(adultdeathrate,"#adult death rate\n"))
        cat(paste(N,"#N\n"))
        cat(paste(numherbivores,"#number of herbivores\n"))
        cat(paste(nl,"#number of loci per herbivore\n"))
        cat(paste(paste(rep(effectsize,nl),collapse=" "),"#effect sizes of metabolites on herbivores\n")) #generalist
        cat(paste(paste(rep(-effectsize,nl),collapse=" "),"#effect sizes of metabolites on herbivores\n")) #specialist
        cat(paste(paste(rep(pi,numherbivores),collapse=" "),"#herbivore occurrence probability\n"))
        cat(paste(paste(rep(0,numherbivores),collapse=" "),"#herbivore introduction times\n"))
        cat("0.0001 #mutation rate\n")
        cat("l #benefit function\n")
        cat("e #cost function\n")
        cat(paste(DB[treatment],"#benefit dominance\n"))
        cat(paste(DC[treatment],"#cost dominance\n"))
        cat(paste(b0,zhalf,senspar,"#benefit parameters\n"))
        cat(paste(c,"#cost parameters\n"))
        cat("m #combine traits\n")
        cat("0.5 #initial allele frequency\n")
        cat("1 #first sampling time\n")
        cat("100 #sampling interval\n")
        cat(paste(N,"#samplesize\n"))
        sink()
        system(paste("./simplesimchemdiv",inputfilename))
    }
}
```
