## Supplemental Code S1 for "How does plant chemodiversity evolve? Testing five hypotheses in one population genetic model": individual.cpp

```
//Copyright 2017, Meike Wittmann

//This file is part of the simulation program used in Wittmann et al. (2017) Segregation lift: How seasonally fluctuating selection can maintain polymorphism at many loci.

//This simulation program is free software: you can redistribute it and/or modify it under the terms of the GNU General Public License as published by the Free Software Foundation, either version 3 of the License, or (at your option) any later version.

//This software is distributed in the hope that it will be useful, but WITHOUT ANY WARRANTY; without even the implied warranty of MERCHANTABILITY or FITNESS FOR A PARTICULAR PURPOSE.  See the GNU General Public License for more details.

//You should have received a copy of the GNU General Public License along with this software.  If not, see <http://www.gnu.org/licenses/>.

#include "individual.hh"
#include <boost/random.hpp>
#include <boost/math/constants/constants.hpp>

extern boost::mt19937 rng;

//initialize static members
int individual::numloci;
int individual::numherbivores;
std::vector<std::vector<double> > individual::alleffectsizes;
double individual::mutationrate;
char individual::benefitfunction;
char individual::costfunction;
double individual::dominance_benefit;
double individual::dominance_cost;
std::vector<double> individual::benefitparameters;
std::vector<double> individual::costparameters;
char individual::combinetraits;

const double pi = boost::math::constants::pi<double>();

individual::individual(const std::vector<int>& mutations1, const std::vector<int>& mutations2)
{
    for (unsigned i=0; i<mutations1.size(); i++){
      (mutations)[mutations1[i]]++;
    }
    for (unsigned i=0; i<mutations2.size(); i++){
      (mutations)[mutations2[i]]++;
    }
}

individual::individual(const std::map<int,int>& m)
  : mutations(m)
{
}

individual::individual(double p0)
{
  boost::binomial_distribution<> binodist(2,p0);
   boost::variate_generator<boost::mt19937&, boost::binomial_distribution<> > copynumber(rng, binodist);
  for (int i=0; i<numloci; i++){
    mutations[i]=copynumber();
  }
}

double individual::fitness(const std::vector<bool>& herbivorepresence){
  double logw = 0;
  //std::map<int,int>::const_iterator end = mutations.end();
  //std::cout << "Genotype: ";
  for (int nh=0; nh< numherbivores; nh++){
    double contribution = 0;
    double benefittraitvalue = 0;
    double costtraitvalue = 0;
    for (int l=0; l<numloci; l++){//all loci contributing to this trait
      //std::cout << mutations[l];
      if (mutations[l]==2){
	benefittraitvalue += alleffectsizes[nh][l];
	costtraitvalue += 1;
      }
      else if (mutations[l]==1){
	benefittraitvalue += dominance_benefit * alleffectsizes[nh][l];
	costtraitvalue += dominance_cost;
      }   
    }
    
    if (combinetraits == 'm'){
      switch (benefitfunction){
      case 's': //saturating
	if (herbivorepresence[nh]){
	  double b0 = benefitparameters[0];
	  double zhalf = benefitparameters[1];
	  
	  contribution += log(b0 + (1-b0)*benefittraitvalue/(benefittraitvalue+zhalf));
	}
	break;
      case 'l': //logistic; works also if there are attractive effects of metabolites on herbivores
	if (herbivorepresence[nh]){
	  double b0 = benefitparameters[0];
	  double zhalf = benefitparameters[1];
	  double senspar = benefitparameters[2];
	  
	  contribution += log(b0 + (1-b0)/(1+exp(-senspar*(benefittraitvalue-zhalf))));
	}
	break;	
      }

      switch (costfunction){
      case 'e': //exponential
	double c = costparameters[0];
	contribution -= c*costtraitvalue;
	break;
      }
      //std::cout << "(" << contribution << ")";
      logw += contribution;
    }
    else {
      std::cerr << combinetraits << " is not a valid model for fitness across traits." << std::endl;
    }
    //std::cout << ",";
  }
  //std::cout << "log fitness: " << logw << std::endl;
  return logw;
}

std::vector<int>  individual::inherit_mutations() const{
  std::vector<int> passed_mutations;
  boost::bernoulli_distribution<> coindist(0.5);
  boost::variate_generator<boost::mt19937&, boost::bernoulli_distribution<> > faircoin(rng, coindist);
 
  std::map<int,int>::const_iterator end = mutations.end(); 
  for (std::map<int,int>::const_iterator it = mutations.begin(); it != end; ++it) {
    int copynumber = it->second;
    if (copynumber==2 || ((copynumber == 1) && faircoin())){ //faircoin is only evaluated if copynumber is 1
      passed_mutations.push_back(it->first);
    }
  }
    
  //implement mutations
  boost::bernoulli_distribution<> coindist2(mutationrate);
  boost::variate_generator<boost::mt19937&, boost::bernoulli_distribution<> > mutationcoin(rng, coindist2);
  std::vector<bool> mutlocus;
  for (int l=0; l<numloci; l++){
    mutlocus.push_back(mutationcoin());
  }
  unsigned m=0;
  while (m<passed_mutations.size()){
    if (mutlocus[passed_mutations[m]]){
      mutlocus[passed_mutations[m]]=false;
      passed_mutations.erase(passed_mutations.begin()+m);
    }
    else {
      m++;
    }
  }
  for (int l=0; l<numloci; l++){
    if (mutlocus[l]){
      passed_mutations.push_back(l);
    }
  }
 
  return passed_mutations;
}

void individual::reset_mutation(int mutid, int copynumber){
  mutations[mutid]=copynumber;
}

void individual::set_mutation_parameters(int nl, int nh, std::vector<std::vector<double> > esm, double mr, char bf, char cf, double db, double dc, const std::vector<double>& bps, const std::vector<double>& cps, char cbt){
  numloci = nl;
  numherbivores = nh;
  alleffectsizes = esm;
  mutationrate = mr;
  benefitfunction = bf;
  costfunction = cf;
  dominance_benefit = db;
  dominance_cost = dc;
  benefitparameters = bps;
  costparameters = cps;  
  combinetraits = cbt;
}
```
