## Supplemental Code S1 for "How does plant chemodiversity evolve? Testing five hypotheses in one population genetic model": Makefile

```
all: simplesimchemdiv
CXXFLAGS= -O3 -Wall -std=c++11 -static


simplesimchemdiv: simplesimchemdiv.cpp population.o individual.o
population.o: population.cpp population.hh
individual.o: individual.cpp individual.hh

clear: 
	rm *.o simplesimchemdiv
```
