## Supplemental Code S1 for "How does plant chemodiversity evolve? Testing five hypotheses in one population genetic model": population.cpp

```
//Copyright 2017, Meike Wittmann

//This file is part of the simulation program used in Wittmann et al. (2017) Segregation lift: How seasonally fluctuating selection can maintain polymorphism at many loci.

//This simulation program is free software: you can redistribute it and/or modify it under the terms of the GNU General Public License as published by the Free Software Foundation, either version 3 of the License, or (at your option) any later version.

//This software is distributed in the hope that it will be useful, but WITHOUT ANY WARRANTY; without even the implied warranty of MERCHANTABILITY or FITNESS FOR A PARTICULAR PURPOSE.  See the GNU General Public License for more details.

//You should have received a copy of the GNU General Public License along with this software.  If not, see <http://www.gnu.org/licenses/>.

#include "population.hh"
#include "individual.hh"
#include <math.h>
#include <boost/random.hpp>
extern boost::mt19937 rng;


population::population(int N, double initfreq)
  : fitnessvector(N,1), fitnessscale(1)
{
  for (int i=0; i<N; i++){
    individual I1(initfreq);
    inds.push_back(I1);
  }
}

void population::resample(const std::vector<bool>& herbivorepresence, int N, double adultdeathrate){
  std::vector<individual> newinds;
  int Nold = inds.size();
  
  boost::uniform_real<> dist(0,1);
  boost::variate_generator<boost::mt19937&, boost::uniform_real<> > uni(rng, dist);

  //survival of adults
  for (int j=0; j<Nold; j++){
    double luck1 = uni();
    if (luck1 > adultdeathrate){//individual survives to next time step
      newinds.push_back(inds[j]);
    }
  }

  //find out the number of free spaces
  int freespaces = N-newinds.size();
  
  //reproduction
  double totalfitness = 0;
  fitnessscale = inds[0].fitness(herbivorepresence);
  for (int i=0; i<Nold; i++){
    double f = exp(inds[i].fitness(herbivorepresence)-fitnessscale);
    fitnessvector[i] = f;
    totalfitness += f;
  }

  for (int j=0; j<freespaces; j++){
    //pick two parents
    int p1 = -1;
    double cum = 0;
    double luck =  uni()*totalfitness;
    while ((cum <= luck) && (p1 < Nold-1)){
      p1 ++;
      cum += fitnessvector[p1];
    }
    
    int p2 = -1;
    cum = 0;
    luck =  uni()*totalfitness;
    while ((cum <= luck) && (p2 < Nold-1)){
      p2 ++;
      cum += fitnessvector[p2];
    }
    
    individual newkid(inds[p1].inherit_mutations(),inds[p2].inherit_mutations());
    newinds.push_back(newkid);
  }

  inds = newinds;
}


std::vector<int> population::take_sample(int samplesize, int numloci) {
  std::vector<int> mutationcounts(numloci,0);
  for (int i=0; i<samplesize; i++){
    std::map<int,int> individual_sample = inds[i].get_mutations();
    std::map<int,int>::const_iterator end = individual_sample.end(); 
    for (std::map<int,int>::const_iterator it = individual_sample.begin(); it != end; ++it) {
      mutationcounts[it->first] += it->second;
    }
  }
  return(mutationcounts);
}

double population::get_frequency(int mutid){
  int mutcount = 0;
  for (unsigned i=0; i<inds.size(); i++){
    mutcount += inds[i].get_mutationcount(mutid);
  }
  double f = double(mutcount)/double(2*inds.size());
  return f;
}
```
