## Supplemental Code S1 for "How does plant chemodiversity evolve? Testing five hypotheses in one population genetic model": simplesimchemdiv.cpp

```
//Copyright 2017, Meike Wittmann

//This file is part of the simulation program used in Wittmann et al. (2017) Segregation lift: How seasonally fluctuating selection can maintain polymorphism at many loci.

//This simulation program is free software: you can redistribute it and/or modify it under the terms of the GNU General Public License as published by the Free Software Foundation, either version 3 of the License, or (at your option) any later version.

//This software is distributed in the hope that it will be useful, but WITHOUT ANY WARRANTY; without even the implied warranty of MERCHANTABILITY or FITNESS FOR A PARTICULAR PURPOSE.  See the GNU General Public License for more details.

//You should have received a copy of the GNU General Public License along with this software.  If not, see <http://www.gnu.org/licenses/>.

#include "individual.hh"
#include "population.hh"
#include <iostream>
#include <iomanip>
#include <boost/random.hpp>
#include <boost/accumulators/accumulators.hpp>
#include <boost/accumulators/statistics/stats.hpp>
#include <boost/accumulators/statistics/mean.hpp>
#include <boost/accumulators/statistics/variance.hpp>
#include <boost/bind.hpp>
#include <boost/ref.hpp>
#include <ctime>
#include <vector>
#include <algorithm> //for min_element and max_element
#include "math.h"
#include <fstream>
#include "string.h"
#include <stdexcept>

boost::mt19937 rng; //global random number generator, declared extern in all other .cc files that need a random number generator

using namespace boost::accumulators;


int numpolymorphic(const std::vector<int>& counts, int N){
  int numpoly =0;
  for (unsigned i=0; i<counts.size(); i++){
    if ((counts[i]>0) && (counts[i]<2*N)) numpoly ++;
  }
  return numpoly;
}

int main(int argc, char* argv[]) {
  time_t start,end;
  time (&start);

  //Check whether the input file is provided and can be opened
  if( argc != 2 )              // File declared?
    {
      std::cerr << "Use: ./simplesim filename" << std::endl;
      return 1;
    }
  std::ifstream inputfile(argv[1]);   // Create  a file stream
  // and open for reading.
  if( !inputfile )                // Get status.
    {
      std::cerr << "An error occurred when opening the file "
		<< argv[1] << std::endl;
      return 2;
    }

  
  std::string filenamebase;
  inputfile >> filenamebase;
  std::cout << "file name base: " << filenamebase << std::endl;
  inputfile.ignore(std::numeric_limits<int>::max(), '\n');

  unsigned seed;
  inputfile >> seed;
  std::cout << "seed: " << seed << std::endl;
  inputfile.ignore(std::numeric_limits<int>::max(), '\n');
  rng.seed(seed);

  int numreps;
  inputfile >> numreps;
  std::cout << "number of replicates: " << numreps << std::endl;
  inputfile.ignore(std::numeric_limits<int>::max(), '\n');

  int generationsperphase;
  inputfile >> generationsperphase;
  std::cout << "generations per phase: " << generationsperphase << std::endl;
  inputfile.ignore(std::numeric_limits<int>::max(), '\n');

  int numgenerations;
  inputfile >> numgenerations;
  std::cout << "number of generations to simulate: " << numgenerations << std::endl;
  inputfile.ignore(std::numeric_limits<int>::max(), '\n');

  double adultdeathrate;
  inputfile >> adultdeathrate;
  std::cout << "adult death rate: " << adultdeathrate << std::endl;
  inputfile.ignore(std::numeric_limits<int>::max(), '\n');
  
  int N;
  inputfile >> N;
  std::cout << "population size: " << N << std::endl;
  inputfile.ignore(std::numeric_limits<int>::max(), '\n');

  int numherbivores;  
  inputfile >> numherbivores;
  std::cout << "number of herbivores: " << numherbivores << std::endl;
  inputfile.ignore(std::numeric_limits<int>::max(), '\n');

  int n_total;
  inputfile >> n_total;
  std::cout << "number of loci: " << n_total << std::endl;
  inputfile.ignore(std::numeric_limits<int>::max(), '\n');


  std::vector<std::vector<double> > effectsizematrix;
  double e1;
  std::cout << "effect sizes: " << std::endl;
  for (int h=0; h<numherbivores; h++){
    std::cout << "herbivore " << h << ": "; 
    std::vector<double> effectsizes1herbivore;
    for (int l=0; l<n_total; l++){
      inputfile >> e1;
      effectsizes1herbivore.push_back(e1);
      std::cout << e1 << " ";
    }
    effectsizematrix.push_back(effectsizes1herbivore);
    inputfile.ignore(std::numeric_limits<int>::max(), '\n');
    std::cout << std::endl;
  }

  std::vector<double> herbivoreprobabilities;
  double p1;
  std::cout << "herbivore occurrence probabilities: ";
  for (int h=0; h<numherbivores; h++){
    inputfile >> p1;
    herbivoreprobabilities.push_back(p1);
    std::cout << p1 << " ";
  }
  std::cout << std::endl;
  inputfile.ignore(std::numeric_limits<int>::max(), '\n');

  std::vector<int> herbivoreintroductiontimes; //times after which the herbivores are present; 0 means the herbivore is present from the beginning
  int it1;
  std::cout << "herbivore introduction times: ";
  for (int h=0; h<numherbivores; h++){
    inputfile >> it1;
    herbivoreintroductiontimes.push_back(it1);
    std::cout << it1 << " ";
  }
  std::cout << std::endl;
  inputfile.ignore(std::numeric_limits<int>::max(), '\n');

  double mutrate;
  bool deactivate_mutations = false;
  int deactivationtime = 0;
  inputfile >> mutrate;
  std::cout << "mutation rate: " << mutrate << std::endl;
  inputfile.ignore(1);
  bool infothere = (inputfile.peek() != int('#'));
  if (infothere){
    inputfile >> deactivationtime;
    deactivate_mutations = true;
  }
  inputfile.ignore(std::numeric_limits<int>::max(), '\n');
  
  char benefitfunction;
  inputfile >> benefitfunction;
  std::cout << "benefit function " << benefitfunction << std::endl;
  inputfile.ignore(std::numeric_limits<int>::max(), '\n');

  char costfunction;
  inputfile >> costfunction;
  std::cout << "cost function " << costfunction << std::endl;
  inputfile.ignore(std::numeric_limits<int>::max(), '\n');

  double benefit_dominance;
  inputfile >> benefit_dominance;
  std::cout << "benefit dominance: " << benefit_dominance << std::endl;
  inputfile.ignore(std::numeric_limits<int>::max(), '\n');

  double cost_dominance;
  inputfile >> cost_dominance;
  std::cout << "cost dominance: " << cost_dominance << std::endl;
  inputfile.ignore(std::numeric_limits<int>::max(), '\n');

  std::vector<double> benefitparameters;
  double parameter1;
  inputfile >> parameter1;
  benefitparameters.push_back(parameter1);
  std::cout << "benefit parameters: " << parameter1 << " ";
  inputfile.ignore(1);
  infothere = true;
  while (infothere){
    infothere = (inputfile.peek() != int('#'));
    if (infothere){
      inputfile >> parameter1;
      inputfile.ignore(1);
      benefitparameters.push_back(parameter1);
      std::cout << parameter1 << " ";
    }
  }
  std::cout << std::endl;
  inputfile.ignore(std::numeric_limits<int>::max(), '\n');

  
  std::vector<double> costparameters;
  inputfile >> parameter1;
  costparameters.push_back(parameter1);
  std::cout << "cost parameters: " << parameter1 << " ";
  inputfile.ignore(1);
  infothere = true;
  while (infothere){
    infothere = (inputfile.peek() != int('#'));
    if (infothere){
      inputfile >> parameter1;
      inputfile.ignore(1);
      costparameters.push_back(parameter1);
      std::cout << parameter1 << " ";
    }
  }
  std::cout << std::endl;
  inputfile.ignore(std::numeric_limits<int>::max(), '\n');
  char combinetraits;
  inputfile >> combinetraits;
  std::cout << "function used to combine fitness across traits: " << combinetraits << std::endl;
  inputfile.ignore(std::numeric_limits<int>::max(), '\n');

  double initfreq;
  inputfile >> initfreq;
  std::cout << "Initial allele frequency " << initfreq << std::endl;
  inputfile.ignore(std::numeric_limits<int>::max(), '\n');

  int firstsamplingtime;
  inputfile >> firstsamplingtime;
  std::cout << "first sampling time: " << firstsamplingtime << std::endl;
  inputfile.ignore(std::numeric_limits<int>::max(), '\n');

  int samplinginterval;
  inputfile >> samplinginterval;
  std::cout << "samplinginterval: " << samplinginterval << std::endl;
  inputfile.ignore(std::numeric_limits<int>::max(), '\n');

  int samplesize;
  inputfile >> samplesize;
  std::cout << "samplesize: " << samplesize << std::endl;
  inputfile.ignore(std::numeric_limits<int>::max(), '\n');

  std::string mutationcountfilename = filenamebase + "_mutationcounts.txt";
  std::ofstream mutationcountfile(mutationcountfilename.c_str(),std::ios::trunc); //trunc deletes previous file content

  std::string fitnessfilename = filenamebase + "_fitness.txt";
  std::ofstream fitnessfile(fitnessfilename.c_str(),std::ios::trunc); //trunc deletes previous file content


  boost::uniform_real<> dist(0,1);
  boost::variate_generator<boost::mt19937&, boost::uniform_real<> > uni(rng, dist);

  for (int r=0; r<numreps; r++){
    individual::set_mutation_parameters(n_total,numherbivores,effectsizematrix,mutrate,benefitfunction,costfunction,benefit_dominance,cost_dominance,benefitparameters,costparameters,combinetraits);

    population P(N,initfreq);
    int timesincelastsampling = 0;
    int timesincelastchange = generationsperphase;//so that new herbivore occurrences will be drawn at the beginning
    std::vector<bool> herbivorepresence(numherbivores,false);
    std::vector<std::vector<int> >  sample_collection;
    std::vector<std::vector<double> > fitnessvector_collection;

    for (int time=0; time<numgenerations; time++){

      std::cout << time << std::endl;
      
      //Deactivate mutations if necessary
      if (deactivate_mutations && (time == deactivationtime)){
	mutrate=0;
	individual::set_mutation_parameters(n_total,numherbivores,effectsizematrix,mutrate,benefitfunction,costfunction,benefit_dominance,cost_dominance,benefitparameters,costparameters,combinetraits);
      }

      if (timesincelastchange == generationsperphase){
	//for each herbivore independently determine whether it will be present in the coming phase
	for (int hi=0; hi<numherbivores; hi++){
	  if (time < herbivoreintroductiontimes[hi]){
	    herbivorepresence[hi] = false;
	  }
	  else {
	    double uni1 = uni();
	    //std::cout << uni1 << " " << herbivoreprobabilities[hi] << std::endl;
	    herbivorepresence[hi] = (uni1 < herbivoreprobabilities[hi]);
	  }
	  std::cout << herbivorepresence[hi] << " ";
	}
	std::cout << std::endl;
	timesincelastchange=0;
      }
      timesincelastchange ++;

      timesincelastsampling ++;
      
      P.resample(herbivorepresence,N,adultdeathrate);//individuals reproduce according to fitness, inherit their genetic material, and offspring can mutate

      if ((time >= firstsamplingtime) && (timesincelastsampling >= samplinginterval)){
	std::vector<int> mutsample = P.take_sample(samplesize,n_total);
        
	sample_collection.push_back(mutsample);

	//get fitness statistics
	std::vector<double> F = P.get_fitnessvector();
	double fitnessscale = P.get_fitnessscale();
	accumulator_set<double, stats<tag::mean, tag::variance> > acc;
	for_each(F.begin(), F.end(), boost::bind<void>(boost::ref(acc), _1));
	std::vector<double> fitnessstats;
	fitnessstats.push_back(mean(acc));
	fitnessstats.push_back(sqrt(variance(acc)));
	fitnessstats.push_back(fitnessscale);
	fitnessvector_collection.push_back(fitnessstats);
	
	timesincelastsampling = 0;
      }
      
      
    }

    //write collected samples to file
    for (unsigned samplenum=0; samplenum<sample_collection.size(); samplenum++){
      mutationcountfile << r << " " << (samplenum+1)*samplinginterval;
      for (unsigned j=0; j<sample_collection[samplenum].size(); j++){
	mutationcountfile << " " << sample_collection[samplenum][j];
      }
      mutationcountfile << std::endl;

      fitnessfile  << r << " " << (samplenum+1)*samplinginterval;
      for (unsigned j=0; j<fitnessvector_collection[samplenum].size(); j++){
	fitnessfile << " " << fitnessvector_collection[samplenum][j];
      }
      fitnessfile << std::endl;
    }
  }

  time (&end);
  double dif = difftime (end,start);
  std::cout << "time: " << dif << " seconds." << std::endl;

  return 0;
}
```
